# Measuring the functional complexity of nanoscale connectomes: polarity matters

**DOI:** 10.1101/2024.09.24.614724

**Authors:** Qingyang Wang, Nadine Randel, Yijie Yin, Cameron Shand, Amy Strange, Michael Winding, Albert Cardona, Marta Zlatic, Joshua T. Vogelstein, Carey E. Priebe

## Abstract

The emerging electron microscopy connectome datasets provides connectivity maps of the brains at single cell resolution, enabling us to estimate various network statistics, such as connectedness. We desire the ability to assess how the functional complexity of these networks depends on these network statistics. To this end, we developed an analysis pipeline and a statistic, XORness, which quantifies the functional complexity of these networks with varying network statistics. We illustrate that actual connectomes have high XORness, as do generated connectomes with the same network statistics, suggesting a normative role for functional complexity in guiding the evolution of connectomes, and providing clues to guide the development of artificial neural networks.

## 1 Introduction

How do brains support complex behaviors? Despite the significant advancements in neuroscience, it remains challenging to identify the fundamental wiring rules or structural properties that are necessary for complex functionalities.

Part of the difficulty lies in expanding the repertoire of structural properties to be studied. Although graph and network theory offer many valuable insights into which network properties might be critical [1, 2], many of these properties are hard to estimate without large-scale, synapse-level connectivity data. Even basic graph metrics, such as in-degree (the number of pre-synaptic neurons), cannot be accurately determined without a complete synapse-level connectivity map. The recent emergence of nanoscale electron microscopy (EM) connectomes solve this problem, enabling analysis at an unprecedented level [3–11].

Another challenge arises from the generality of the concept of functional complexity, rendering it difficult to define or quantify without relying on specific behaviors or tasks. Statistical theories, such as Vapnik–Chervonenkis dimension and Rademacher Complexity [12] offer analogous means of defining functional complexity; yet, they depend on exhaustive metrics that are difficult to translate into experimentally testable measurements. To address this, we propose a functional complexity measurement that is task-agnostic, learning-independent, and experimentally testable.

The brain structure property we focus on is the Excitatory-Inhibitory (E-I) neuron ratio. Although basic, the E-I ratio is highly conserved across species. Yet, its functional significance remain poorly understood. Across a wide range of species—from *Drosophila* [4, 13], rodents [14–20], cats [21–23], to non-human primates [24], and humans [25]—excitatory neurons are the majority of neurons (∼70-90%, Figure 6A). This Excitatory-Inhibitory (E-I) ratio is also conserved across different functional regions of the neocortex [14, 21, 24, 25]. What is the necessity of having so many excitatory neurons? Our results offer a clue.

By sampling many thousands of recurrent neural networks (RNNs) constrained by the larval and adult *Drosophila* EM connectome [3] and quantifying their functional complexity, we demonstrate that the optimal E-I ratio that maximizes functional complexity favors a high proportion of excitatory neurons: the top 1% networks have an average of *>*70% excitatory neurons. Furthermore, we demonstrate that the differential connectedness between excitatory and inhibitory neurons—inhibitory neurons tend to have more connections—also interplays with E-I ratio to optimize the overall functional complexity of networks. The E-I ratio and connectedness distribution of the top functional complexity simulations closely mirrors that of real brains. In the end, we close the loop by applying the insights extracted from EM connectome to solve a challenging scenario in artificial intelligence.

## 2 Results

### 2.1 Measuring functional complexity based on nonlinearity

We give the full detail of the functional complexity measurement in Section 4; here, we focus on building the intuition and reasoning behind the measurement. Consider an illustrative example where an animal has to classify toxic versus edible leaves (a binary classification problem). In *world alpha*, toxic leaves are always red regardless of shape (Figure 1B left two illustrations). In *world beta*, toxic leaves (class A solid) could be either green straight or red curved; edible leaves (class B dashed) could be either green curved or red straight. *World beta* requires solving more complicated function, because it cannot be solved with a linear function.

**Figure 1:**
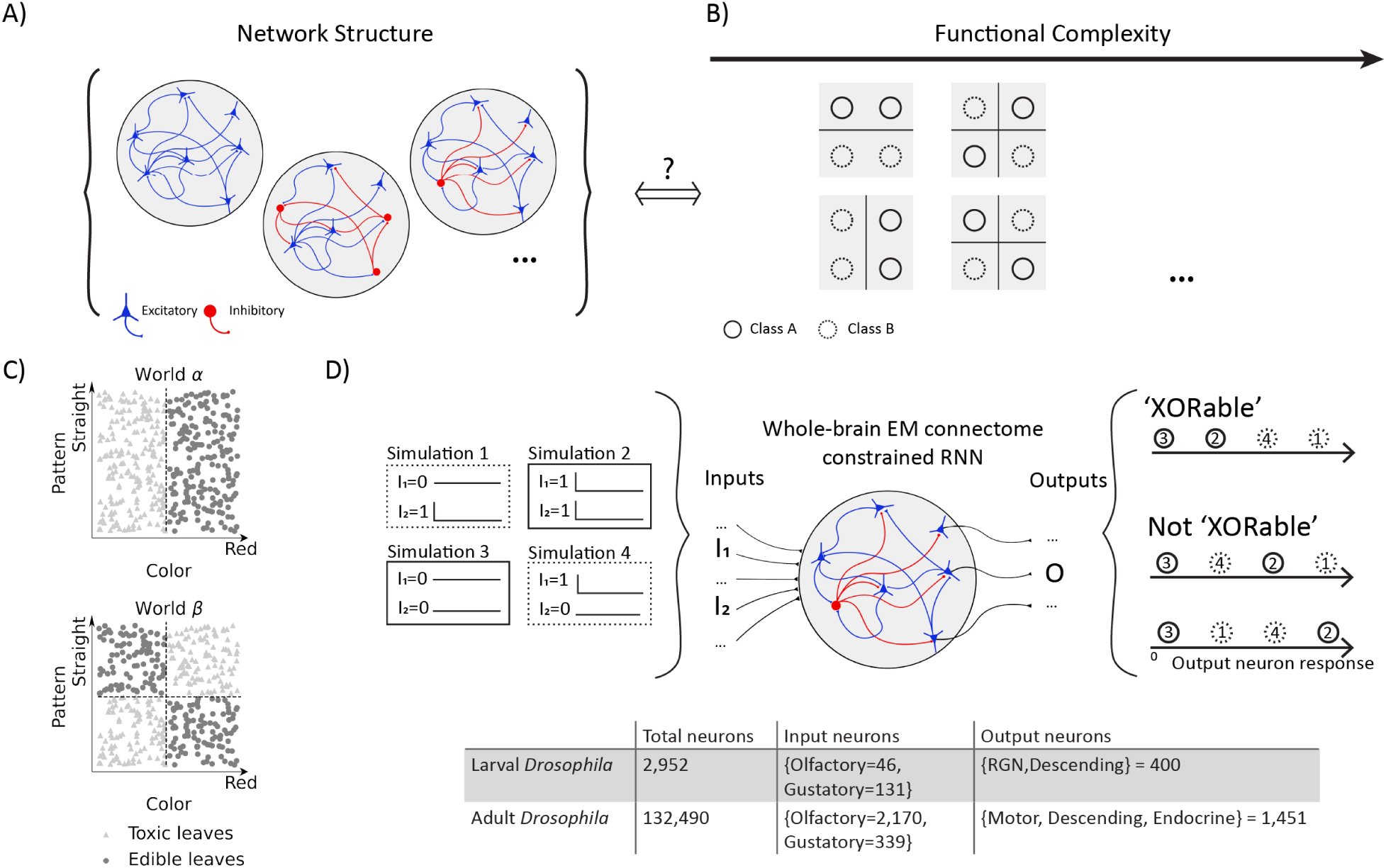
Proposed methodology to measure functional complexity of a given network. A) Illustration on relating network structure, specifically E-I composition, to the network’s functional complexity. B) XOR is a binary classification that takes two inputs, with the desired output being the two scenarios where only one of the two inputs is activated are classified into the same class (second and forth quadrant, dashed circles). Whether a subnetwork is ‘XORable’ serves as the building block of the functional complexity measurement and it can be tested by four independent input stimulation pairs. C) Given an EM-constrained larval *Drosophila* whole-brain RNN (2952 neurons in total, E-I composition sampled as detailed in Section 4), we iterate through the subnetworks {*I*1*, I*2*, O*} where each subnetwork is defined by {*I*1*, I*2} as two input neurons and *O* as the output. Exhaustively, we measure through all olfactory and gustatory neurons as inputs and all 400 output neurons (RGN and descending neurons). Whether a subnetwork {*I*1*, I*2*, O*} is XORable is tested by the following procedure: four independent simulations are run (illustrated on the left and in panel C). The four simulation inputs correspond to {(0, 1), (1, 1), (0, 0), (1, 0)} respectively, where 1 means the input neuron is activated. XORable corresponds to the situation where simulations 1 and 4 elicit output response that is linearly separable from simulations 2 and 3. In another word, the relative order of 1 and 4 should be larger than 2 and 3. This is illustrated on the right along with counter-examples.

*World beta* is actually a description of XOR, a binary classification problem that involves two dimensional input and a binary output. In fact, any binary classification task that is dependent on both features is an XOR task [26]. Multiple animal behavior studies have shown that XOR is more challenging for humans and monkeys to learn compared to linearly solvable tasks [26–29]. Even in machine learning research, XOR has played a pivotal role in revealing the functional complexity of networks: the first AI winter started when Minsky and Papert pointed out perceptrons cannot solve XOR [30]. XOR has also revealed the functional significance of specific network structure: in prior work we proved that networks with non-decreasing activation functions (e.g. ReLU) and without any negative weights (equivalent to inhibitory connections) cannot solve XOR and thus are not universal approximators^1^ [34].

Given a network with *n* neurons, out of which |I| are input neurons and |O| are output neurons, we measure its functional complexity by considering how many subnetworks can readily solve XOR, termed XORable subnetworks. The more XORable subnetworks, the more nonlinear input associations the network is able to form and the more complex decision boundaries the network can represent. A subnetwork is defined by the input-output combination {*I*_1_*, I*_2_*, O*}, which implicitly considers all the possible paths connecting the two input neurons to the output neuron within certain number of time steps. The procedure for deciding whether a subnetwork is XORable is illustrated in Figure 1D. Briefly, four independent simulations are run where the two input neurons are combinatorially activated at time 0 (left panel), corresponding to the four quadrants of the 2D input space (Figure 1C). The subnetwork {*I*_1_*, I*_2_*, O*} is XORable when the output activity of the four simulations follow a particular order: the two simulations where both input neurons are simultaneously activated or not activated (solid borders in Figure 1C-D) should elicit output response that is linearly separable from the simulations where only one of the input neurons was activated. We iterate through all concerned subnetworks with the above procedure to obtain the functional complexity measurement.

We did not involve any parameter optimization in the functional complexity measurement: the connectome already encapsulates knowledge from evolution and life experiences, and our goal is to quantify the functional complexity on connectivity structures that is naturally present.

### 2.2 EM-constrained RNNs of varied E-I configurations show varied functional complexity

The recurrent neural network (RNN) (details in Section 4) is constrained by the larva *Drosophila* EM connectome [3]. There are in total 2952 neurons that form a connected graph (one single connected component), 400 of which are output neurons (RGNs and descending neurons) and 480 are input neurons. We will focus on olfactory (46) and gustatory (131) neurons as input throughout this paper. The neurons’ firing rates **r** at time *t* are modeled as

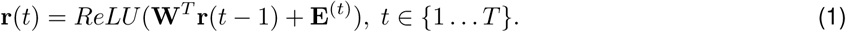

**E** is the external stimuli in Figure 1D. The connectivity from pre-synaptic neurons to post-synaptic neurons, **W**, is the element-wise product of two components: 1) the strength of connections **M**, which is given by the synaptic count from the EM dataset (axon-dendrite synapses in main text; all-all connection results in the supplementary); and 2) the excitatory/inhibitory identity ***α*** of each neuron which is randomly sampled.

The 2D XOR patterns were presented to the model by either stimulating two olfactory neurons (Figure 2 second column), two gustatory neurons (third column), or one olfactory paired with one gustatory neuron (rightmost column). We first varied E-I ratio by uniform Bernoulli sampling (horizontal lines in left column), i.e. all neurons have the same probability *p* of being excitatory. As a sanity check of our method, consider the extreme case where all neurons are excitatory: previous theoretical results predicted no XORable subnetwork exists in this scenario [34]. This is exactly what we observed, when E/(E+I)=1 we observed zero functional complexity across all panels. The optimal E-I ratio that maximized functional complexity fell between ∼50-60% excitatory neurons. In real larval brains, the ratio is closer to 67% [13].

**Figure 2:**
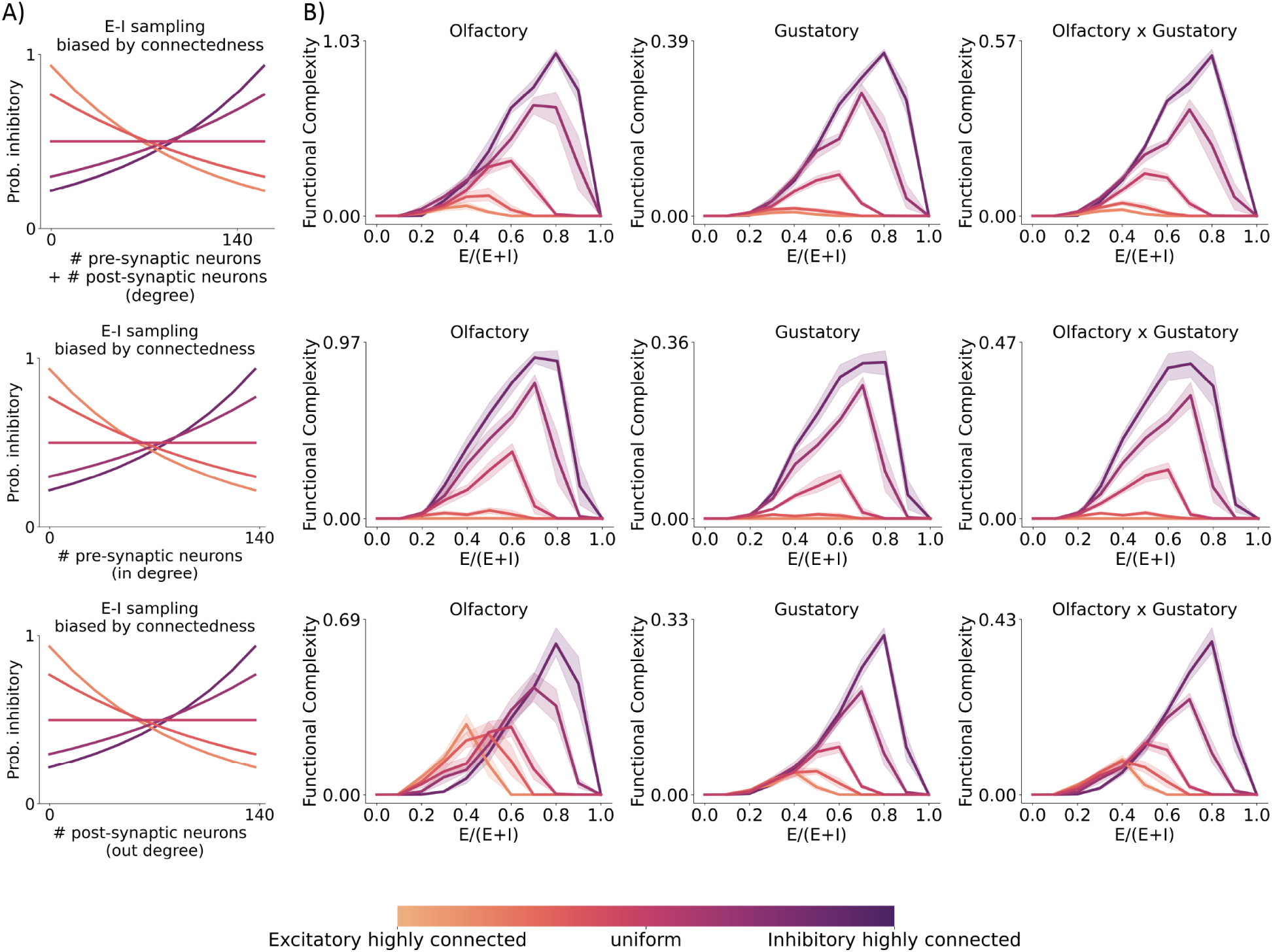
The optimal E-I ratio that maximizes functional complexity is dependent on the connection degree of inhibitory neurons. A) E-I probability as a function of its connectedness, quantified as either in degree (number of pre-synaptic neurons), out degree (number of post-synaptic neurons), or degree (in+out). For each neuron, the probability for it being sampled as inhibitory depends on its connectedness. All colored curves result in a target E-I ratio of 0.5 but with different extent of biasing the inhibitory population to be highly connected (noted by the colors given in the bottom colorbar). Darker colors = inhibitory neurons are more highly connected; Lighter colors = excitatory neurons are more highly connected. B) The resultant functional complexity with E-I identities sampled at various E-I ratio levels (x-axis) and various connectedness bias levels (colors). Shaded area mark the 95% confidence interval (n=10). When the E-I identity is uniformly sampled across all neurons (horizontal line in A and the curves of middle color in B), the optimal E-I ratio that maximizes functional complexity is ∼60% excitatory, across sensory modalities. However, the maximum level of functional complexity reached by uniform sampling is sub-optimal; more optimal E-I compositions require the sampling procedure to bias the inhibitory neurons to be highly connected (darker colors). In contrast, when the E-I sampling procedure shifts the excitatory neurons to be highly connected (lighter colors), it generally results in lower functional complexity.

This discrepancy prompted us to think maybe considering E-I ratio alone was insufficient, therefore we designed a procedure to sample the E-I identities as a degree dependent function (Figure 2A): for each neuron, the probability of it being sampled as inhibitory is dependent on its connectedness (details in Section 4). Ubiquitous evidence from the mouse cortex showed that inhibitory neurons are broadly tuned integrating diverse input [35–37] and sending out inhibition broadly [38, 39]. Therefore we quantified the connectedness of neurons in three ways: in degree (# pre-synaptic neurons) quantifying integration diversity, out degree (# post-synaptic neurons) quantifying output diversity, degree (# pre-synaptic neurons + # post-synaptic neurons) combining both. With the connectedness-dependent sampling procedure, we systematically varied the network structure from having highly connected excitatory neurons (lighter colors) to having highly connected inhibitory neurons (darker colors). Intriguingly, as the network structure shifted from highly connected excitatory neurons to highly connected inhibitory neurons (light → dark), the optimal E-I ratio peak gradually shifted from 30-40% to 80% (Figure 2B). Furthermore, the overall optimality is achieved when the inhibitory neurons are highly connected and at 80% E-I ratio. These two observations hold across input sensory modalities (columns), and for both integration connectedness and output connectedness (rows).

These results suggest to achieve high functional complexity, it is desirable to have a good fraction of excitatory neurons, with inhibitory neurons receive broad input and output broadly. Although these results are based on axon-dendrite synaptic counts, they still hold when the connection magnitudes are given by all synapse types, i.e. axo-dendritic, axo-axonic, dendro-dendritic, and dendro-axonic (Figure A.3).

### 2.3 The RNNs with highest functional complexity match the E-I ratio and degree distribution of real brains

Does our observation from the previous sections generalize? That is, is it generally true that networks with over-abundance of excitatory neurons and highly-connected inhibitory neurons enjoy higher functional complexity? Next we quantified the functional complexity of an extensive set of EM connectome-constrained networks of various E-I configurations sampled by various forms of connectedness-dependent sampling procedures, including ones where the trend is not monotonic, i.e. neurons of mid-level connectedness having high probability of being inhibitory/excitatory (details in Section 4). In total, we performed 818 different types of sampling procedures according to different connectedness-dependent E-I probability functions, each type with 10 different samples (Figure 3). Notably, networks of higher functional complexity (lighter color) not only have an over-abundance of excitatory neuron population (x-axis > 0.5, right to vertical gray line), they also have more highly connected inhibitory population compared to the excitatory population (y-axis > 0, above horizontal gray line). The degree difference between inhibitory and excitatory population is evident when considering input integration (middle column, more in Supplementary Figure A.10,A.12), output breadth (right column, more in Supplementary Figure A.14,A.16), or combined (left column, more in Supplementary Figure A.8). They also hold across input from different sensory modalities (Supplementary Figure A.6, A.7, A.9, A.11, A.13, A.15).

**Figure 3:**
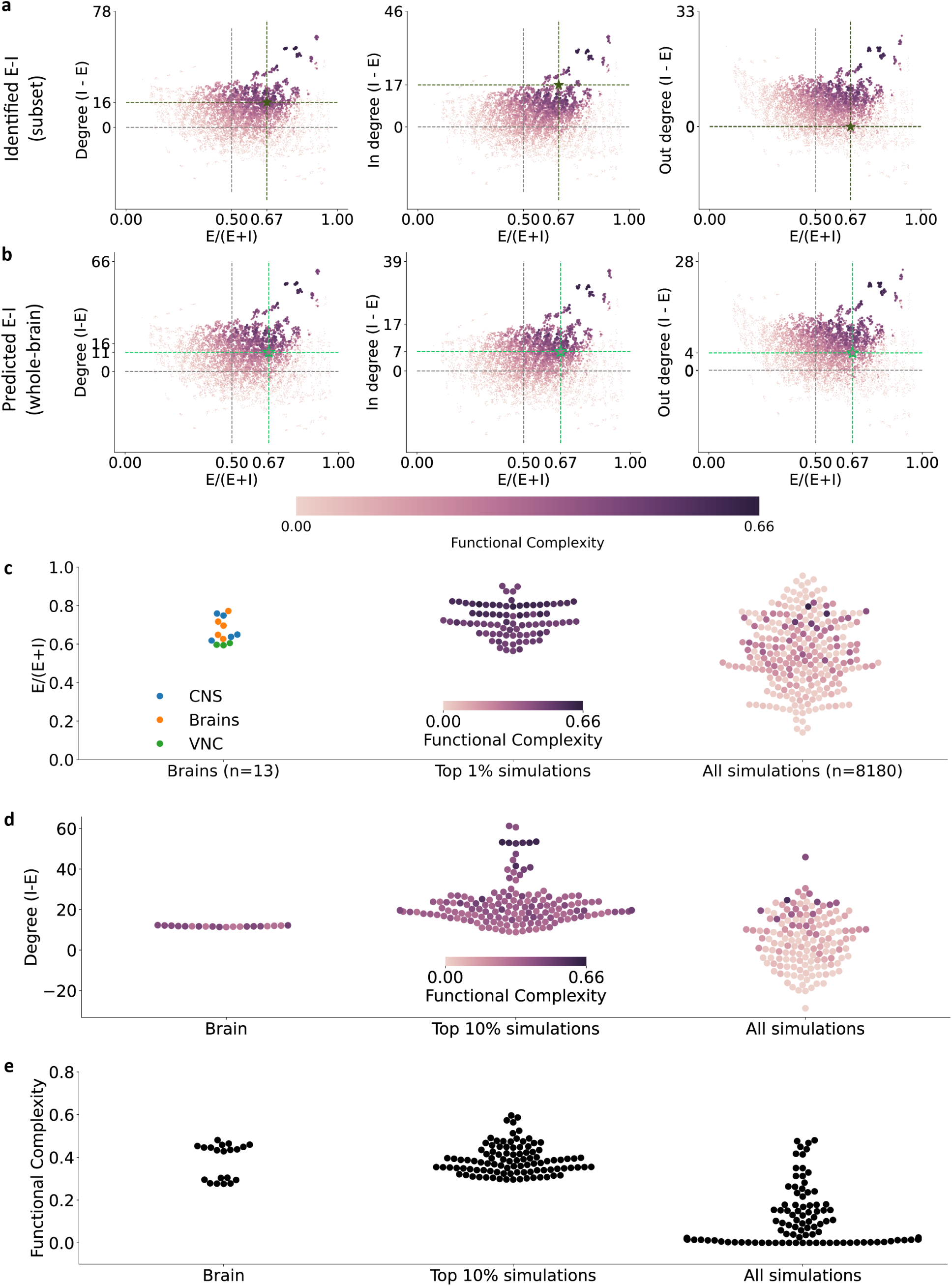
The top simulations of highest functional complexity match the E-I ratio distribution observed in real larval *Drosophila* brains. A-B) 8180 different E-I compositions are drawn from 818 different connectedness-dependent E-I probability functions (10 samples each). Networks of higher functional complexity (lighter color) lie on the upper right quadrant where the average connectedness of inhibitory neurons are higher than excitatory neurons (y-axis > 0, above horizontal gray line) and there is an over-abundance of excitatory neurons (x-axis > 0.5, right to vertical gray line). Left to right: differential connectedness of inhibitory and excitatory neurons are measured by degree (in+out), out degree, and in degree respectively. The green stars mark real brain measurements: top (A) x-axis given by scRNA-seq measurements [13] and y-axis given by neurons in the EM connectome with known E-I identities (unpublished data); bottom (B) given by all neurons in the EM connectome with predicted E-I identities (unpublished data) C) Left: E-I ratios of 13 larval *Drosophila* obtained via scRNA-seq, colored by the source region; Middle left: the E-I ratios of top 13 simulations obtained via averaging across the 10 samples of each connectedness-dependent E-I probability function group (details see Supplementary Figure A.17); Middle right: the E-I ratios of top 1% simulations; Right: E-I ratios of all simulations. The mean E-I ratio of the top 13 group-averages and top 1% simulations are *not* statistical significantly different from the 13 brain E-I ratios (two-sided two-sample t-test, p-val: 0.98, 0.51 respectively). Multiple comparisons corrected by Dunnett’s. D) Same as C except for the differential connectedness of inhibitory and excitatory neurons, i.e. Degree (I-E), the brain data is based on EM connectome E-I predictions. E) Same as C except for the functional complexity.

Do the network structural predictions based on functional complexity match to real brain observations? Yes. The real larva measurements (green stars in Figure 3A,B) fall within the region of networks with high functional complexity. In terms of the E-I ratio prediction, the top 1% networks of highest functional complexity have on average 72% of excitatory neurons. The distribution of the E-I ratio of the highest sampled networks closely aligns with the real brain E-I ratios obtained by scRNA-seq [13](green stars x-value in Figure 3A and first column in C). This is true regardless of the exact criterion for selecting the *top* simulations (Figure 3C): the result holds when comparing brain data to top 1% simulations (p-val 0.98, Dunnett’s [40]); it also holds when comparing to the top 13 probability function group averaged E-I ratios (p-val 0.40, Dunnett’s). When comparing the real brain E-I ratio distribution with all simulations, the means are statistically significantly different (p-val: 0.03, Dunnett’s). The top functional complexity simulations’ match to real brain E-I ratios not only hold for networks sampled by degree (in+out), but also hold for networks sampled by in degree (Figure A.10C) as well as out degree (Figure A.14C); the match also holds when the connection weight is given by synapse counts of all synapse types (Figure A.8C, A.12C, A.16C).

To check if the inhibitory neurons tend to be more highly connected in real brains, we need to obtain the E-I identities of the whole-brain EM connectome. In another study (unpublished data), we identified the neurotransmitters for a subset of neurons (170 acetylcholine as excitatory neurons, 156 GABAergic and 26 glutamatergic as inhibitory neurons), marked by the green stars in Figure 3A . Since the selection of the subset is not representative of the whole-brain E-I ratio, we marked the x-axis by scRNA-seq data. Although the subset of neurons with known E-I identities are not selected randomly, their connection properties still exhibit trend matching predicted networks of higher functional complexity (upper right quadrant). In a separate study, we used these known neurotransmitter identities to predict for the rest of the neurons via machine learning method [41], identifying 1917 acetylcholine as excitatory neurons, 160 GABAergic and 857 glutamatergic as inhibitory neurons (unpublished data, the model’s accuracy was 98% for acetylcholine, 59% for GABAergic, and 78% for glutamate expressing neurons). The predicted whole-brain E-I identities are marked by the green stars in Figure 3B, with the color inside the star indicating the measured functional complexity based on the predicted whole-brain E-I identities. Again, the whole-brain observations match to the predicted networks of higher functional complexity (Figure 3D). To account for potential E-I prediction errors, we shuffled the E-I identities of neurons with low prediction confidence (second column in Figure 3D) and did not find it to complicate the conclusion.

Given the whole-brain E-I predictions, we also measured its functional complexity as well as for the low-prediction-confidence shuffled networks (Figure 3E) - they all match to the functional complexity range of the top 10% simulations. These suggest real larval *Drosophila* brain indeed enjoy high functional complexity.

Of note, the functional complexity measurement that we propose is the key in revealing these patterns — a conventional norm-based method such as the participation ratio and spectral norm fail to do so (Supplementary Figure A.5).

Collectively, our extensive simulation results point out two important network structure properties in supporting high functional complexity: 1) over-abundance of excitatory neurons; and 2) highly connected inhibitory neurons. Furthermore, such structural properties selected by functional complexity match to real brain observations.

### 2.4 Results generalize across developmental stages to adult *Drosophila*

To test if our results generalize across developmental stages, we performed the same functional complexity analysis in the adult *Drosophila* whole-brain EM connectome (FlyWire [4]). The FlyWire dataset has 132,490 neurons in total, with 2,170 olfactory neurons, 339 gustatory neurons, and 1,451 output neurons (motor, descending, and endocrine). The adult whole-brain network is orders of magnitudes larger than the larval EM connectome, therefore sub-sampling was adopted to accommodate for the computation requirements (details in Section 4).

We sampled 703 different E-I compositions from 235 different connectedness-dependent E-I probability functions (Figure 4A-B). Although the sampling procedure is carried out in a smaller scale compared to what we did for the larvae, the resultant network structures still span the full feature space under investigation, i.e. E-I ratio and differential connectedness between inhibitory and excitatory populations. The results in adult *Drosophila* show striking similarity to what was observed in the larvae: First, across different input sensory modalities (Figure 4A), higher functional complexity (lighter color) for RNNs constrained by the adult *Drosophila* connectome is associated with both over-abundance of excitatory neuron population (x-axis > 0.5) as well as highly connected inhibitory population compared to the excitatory population (y-axis > 0). Second, as in the larvae, we observe a preference for more highly connected inhibitory neurons (Figure 4B) when we consider the input integration (middle), output breadth (right) and combined (left).

**Figure 4:**
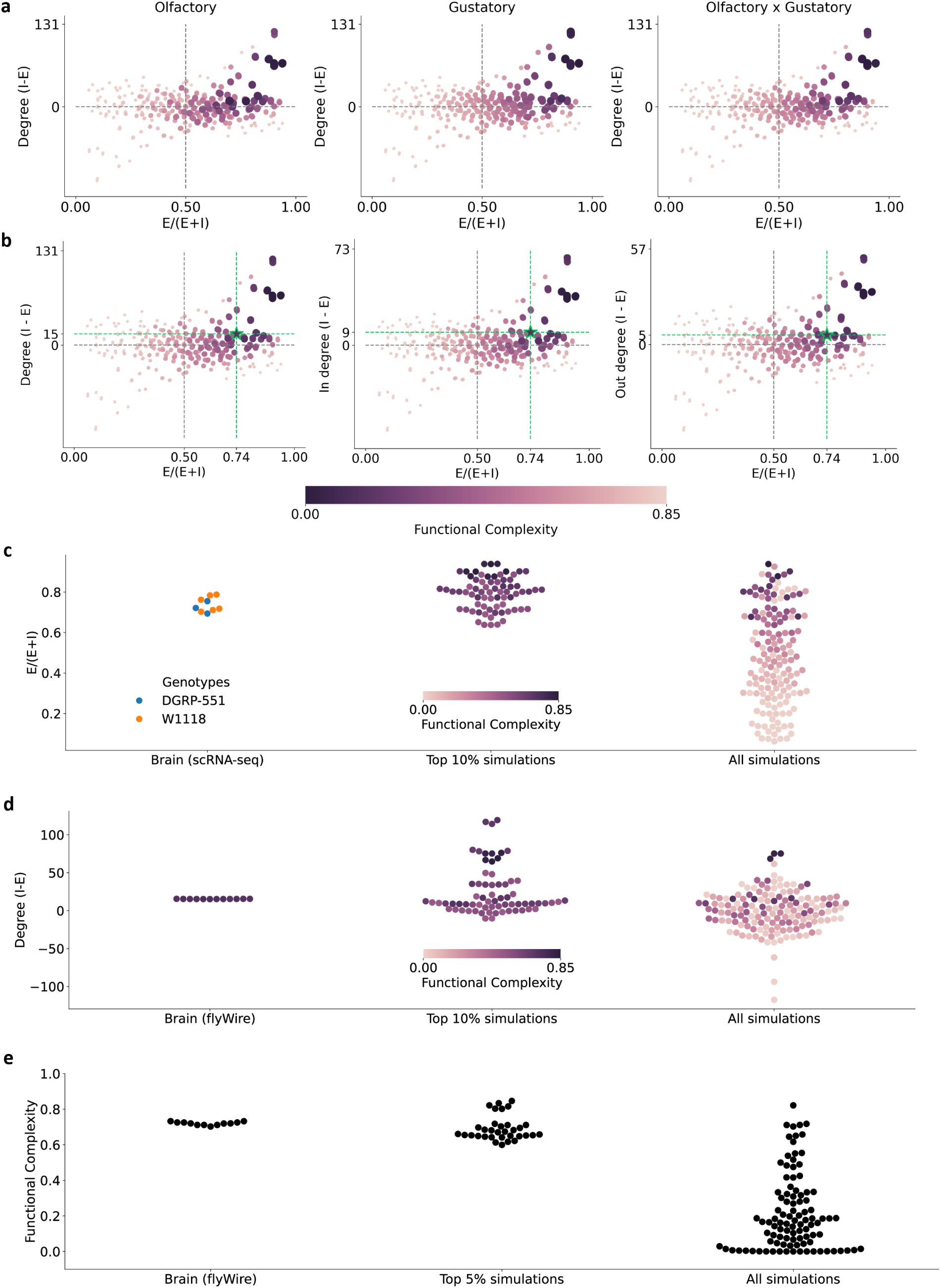
The results generalize across developmental stages to adult *Drosophila*. A-B) 703 different E-I compositions are drawn from 235 different connectedness-dependent E-I probability functions (3 samples each). As observed for the larvae, networks of higher functional complexity (lighter color) lie on the upper right quadrant where the average connectedness of inhibitory neurons are higher than excitatory neurons (y-axis > 0, above horizontal gray line) and there is an over-abundance of excitatory neurons (x-axis > 0.5, right to vertical gray line). A) Left to right: the observations hold across sensory modalities. B) Left to right: differential connectedness of inhibitory and excitatory neurons are measured by degree (in+out), out degree, and in degree respectively. The green stars mark real brain measurements given by neurons in the EM connectome with predicted E-I identities. C) Left: E-I ratios of 9 adult *Drosophila* (age 6d and 9d) obtained via scRNA-seq, colored by the genotypes (data from [42]); Middle left: E-I ratio based on FlyWire neurotransmitter predictions (each dot is a low-prediction-score shuffle); Middle: the E-I ratios of top 10% simulations; Right: E-I ratios of all simulations. D) Same as C except for the differential connectedness of inhibitory and excitatory neurons, i.e. Degree (I-E), the brain data is based on EM connectome E-I predictions. E) Same as C except for the functional complexity.

Furthermore, such predictions based on highest functional complexity also match to real brain observations (green stars). First, the top 10% simulations with highest functional complexity have on average 79% excitatory neurons which is strikingly similar to the 74% average observed from real brains across two genotypes via scRNA-seq [42] (Figure 4C). To see if in the brain the inhibitory neurons in general tend to be more highly connected than excitatory neurons, we again calculated the population average difference (Figure 4D) and took care of the missing E-I predictions by random assignments (left column): indeed, on average, the inhibitory neurons connect with 15 more pre-synaptic and post-synaptic neurons than excitatory neurons. Furthermore, based on the E-I predictions on the

EM connectomes, the measured functional complexity score of real brains mounts to 0.71 on average, which is strikingly similar to the top 5% simulated networks which mounts to 0.72 on average.

Collectively, our functional complexity analysis on adult *Drosophila* whole-brain EM connectome provide a normative explanation to why across developmental stages, *Drosophila* tend to have over-abundance of excitatory neurons and highly-connected inhibitory neurons.

### 2.5 E-I structures identified from the connectome functional complexity analysis solves a challenging scenario in AI

Our functional complexity analysis on larval and adult *Drosophila* EM connectomes suggest that over-abundance of excitatory neurons and highly-connected inhibitory neurons offer an advantage in representing complex functions. Could we leverage this insight to build AI that learn complex tasks better? We next tested these ideas in sparse networks. We focused on the setting where networks are highly sparse because 1) it has been a challenging setting for AI [43]; 2) adequate sign configurations (E-I configurations) *are* crucial in training sparse networks [44, 45]; and 3) biological networks are sparse, both the larval and adult *Drosophila* whole-brain connectomes are 99% sparse. We trained convolutional neural networks (CNNs) at 99% sparsity on Fashion-MNIST, a pattern recognition benchmark (Supplementary Figure A.19). The E-I configurations and wiring were either unconstrained or biologically constrained. In the unconstrained classical pruning scenario, both signs and wiring can be dynamically updated throughout the training process, enjoyings full flexibility (orange); In the biologically constrained scenarios, we asked the signs to follow Dale’s principle, i.e. all output edges of the same neuron follow the same sign, and tested 3 E-I configuration settings: 1) 80% excitatory neurons with 4 times more highly connected inhibitory neurons (red); 2) 50% excitatory neurons with equally connection-degree E/I populations (blue); 3) 20% excitatory neurons with 4 times more highly connected excitatory neurons (green). These three scenarios are designed to achieve equal number of positive and negative weight entries.

Unconstrained sparse networks indeed have trouble learning the task (Supplementary Figure A.19**a**, orange), especially when the networks are deep (convDepth = 4, orange solid dots show significant inferior performance across the board), and have limited width (left two columns). In these challenging scenarios, CNNs that are biologically constrained to follow Dale’s principle do *not* suffer from inferior performance (Supplementary Figure A.19**a**, red, blue, green). Furthermore, over-abundance of excitatory with highly-connected inhibitory neurons demonstrate an advantage over other E-I configurations in the more challenging settings. These are evident from the fact that 80% excitatory group (red) consistently achieve higher validation accuracy at convergence (higher in Figure 5**a**) and takes less number of iterations to achieve 75% validation accuracy (lower in Figure 5**b**). The computational efficiency advantage is independent of the accuracy cut-off selection (Supplementary Figure **??**). These results demonstrate that the E-I structures identified from our functional complexity analysis on EM connectomes offer solutions to train sparse networks effectively and computational efficiently.

**Figure 5:**
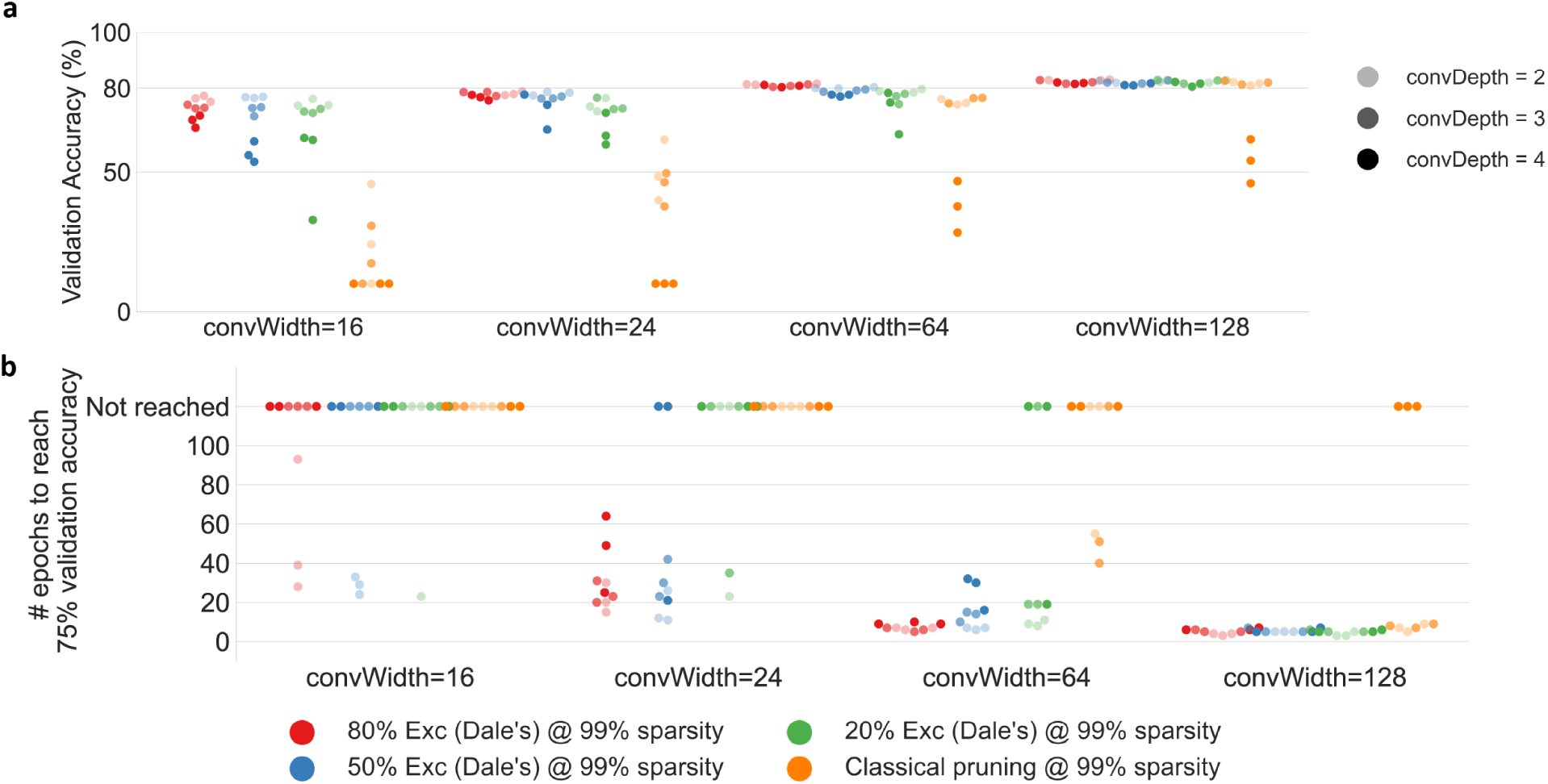
Bio-realistic E-I configurations offer solutions to sparse network training problems. **a** Sparse CNNs constrained by Dale’s rule (red, blue, green) reach higher validation accuracy at convergence then classical pruning procedure (orange), with 80% excitatory setting (red) demonstrating an advantage in difficult settings where the network is narrow and deep (depth=4, width=[16, 24]), i.e. among darkest dots red always dominate. **b** 80% excitatory demonstrates an advantage in computation efficiency as it not only have higher chance of reaching 75% validation accuracy (less dots at the top ‘not reached’ section) but also take comparative or less number of epochs.

**Figure 6:**
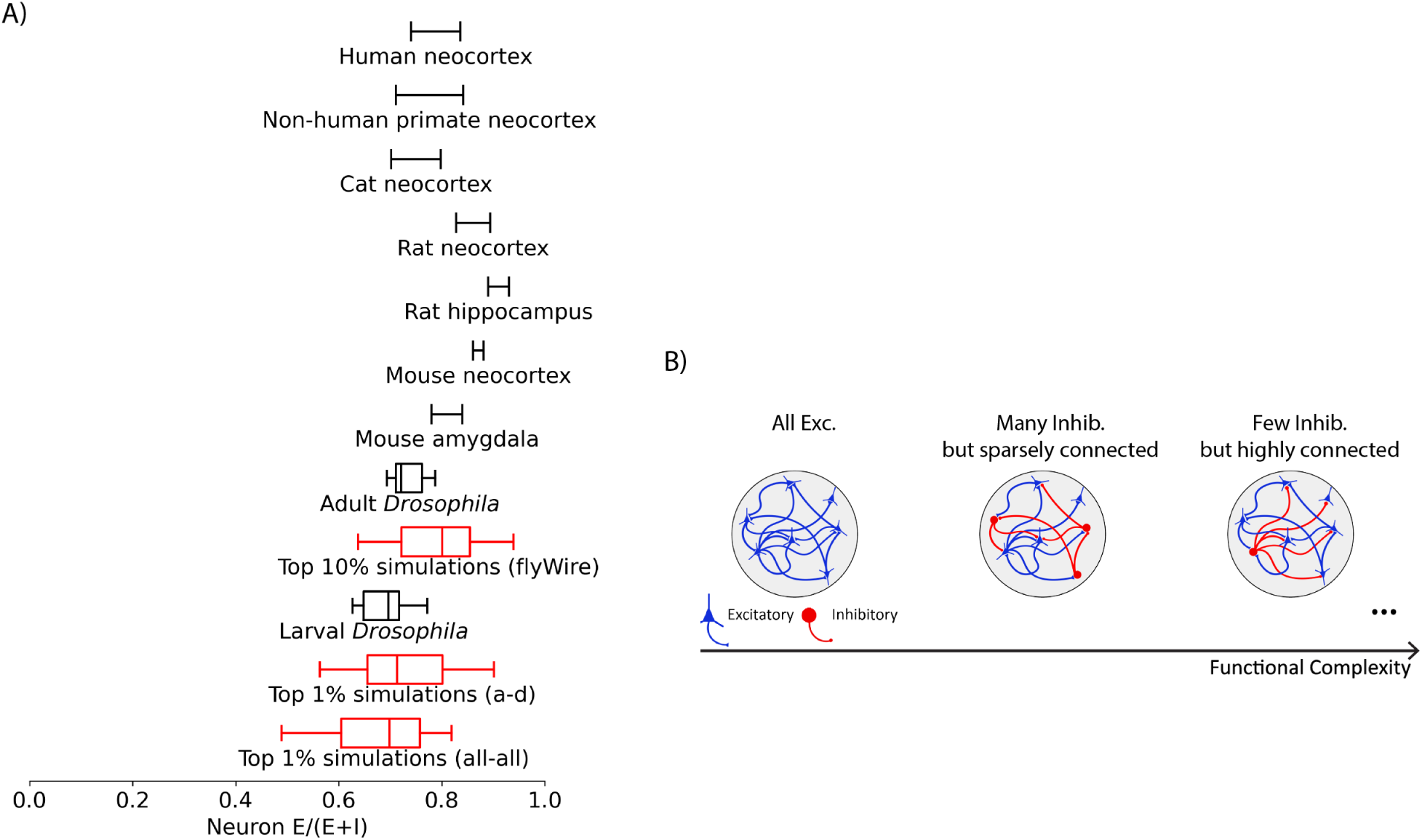
Conceptual illustration on E-I composition and functional complexity. A) The over-abundance of excitatory neurons is observed across a wide range of species (from Drosophila [4, 13, 42] to mouse neocortex and amygdala [14, 15] to rat neocortex [16–18] and hippocampus [19, 20], to cats [21–23] to non-human primates [24] to humans [25]). B) A conceptual illustration on the relationship between E-I composition and functional complexity.

## 3 Discussion

Why do we have so many excitatory neurons? Our answer is: to solve more complex functions. To measure functional complexity of networks, we proposed a normative measurement based on the nonlinearity of subnetwork functions. We leveraged the larva and adult *Drosophila* EM connectome [3] and revealed a key structure property - connectedness of neurons - that alters the functional complexity of the resultant networks. The top E-I configurations with the highest functional complexity nicely matched the E-I ratio and degree distributions observed in real larva and adult*Drosophila* brains. It is worth pointing out that the calculation of degree for all neurons is *not* possible without the availability of the EM connectome data. This emphasizes on the importance of having the entire connectivity map, so that the unexplored but crucial structure properties and their interplay could be explored. We further applied the E-I structures identified from our functional complexity analysis on EM connectomes to offer solutions to train sparse networks effectively and computational efficiently.

The optimal E-I ratio has been previously explored in terms of the capacity on storing binary memory patterns [46]. The predicted optimal E-I ratio is a function of the degree of tuning (coefficient of variation) of the excitatory and inhibitory activities. Consequently, validating it requires knowing both the input statistics and the connectivity pattern. [47] had trained RNNs with sparse connections (out degree following a bio-plausible distribution) to sequentially learn multiple binary classification tasks; consistent with the normative answers provided here, they also found the optimal performing RNNs to have 70% excitatory neurons [47]. These suggest the functional complexity measurement proposed here may have direct implications on networks’ capabilities in multi-task learning.

We deliberately set all bias terms of the RNN, i.e. threshold of the neurons, to be zero. The main reason is we did not want the biases to obscure the effects of the connectivity patterns. It is known that any dynamics’ trajectory can be approximated arbitrarily well by a wide-enough RNN with fixed weights and learnt/well-chosen biases [48]. This means if we choose to learn the bias terms or even just randomly set them, there is a good chance the subnetwork can produce XORable dynamics; but then such observed XORable pattern is not due to the connectivity pattern but because of the non-zero biases, resulting in high false positives when checking for XORable subnetworks. This logic is further backed by the fact that XOR can be solved by a network with only three hidden units and all zero biases [49]; although the 2952 neurons are not fully connected to each other in our case, there is still a high chance non-zero biases will greatly increase the number of XORable subnetworks and obscure the effects from the connectivity pattern. A dedicated separate study is required to carefully delineate the interplay between biases and connectivity pattern in terms of their effects on functional complexity.

The inspiring thread of work on manifold capacity touches upon the concept of representation complexity in classification settings [50–53]; however, the definition of complexity in manifold capacity fundamentally differ from the proposed functional complexity measurement, leading the two methods targeting completely different questions. Manifold capacity targets neuronal activity: essentially, a population of neurons has higher capacity when the desired output can be more readily linearly read-out for the pre-specified task output, leaving the discussion on the non-linearity part of the transformation implicit. In contrast, our functional complexity measurement directly measures the non-linearity of the network. Although the latest development incorporated non-linear classifications into the framework, it relies on context-dependent learning and the resultant capacity measurement is dependent on the definition of context [53]. In sum, manifold capacity focuses on neural representations and their linear readout; ours focuses on directly quantifying the non-linearity of networks.

In statistics and mathematics, some possible choices of quantifying the concept of functional complexity include Rademacher Complexity [54] and VC-dimension [55], which essentially measures the richness of a function class. However, neither can be directly applied because they quantify complexity of a function class, and we are interested in complexity of a particular function: the actual weights of a particular network. The proposed functional complexity measurement relates to VC-dimension in the sense that it considers the hardest problems (XOR like) out of the entire function set considered in VC-dimension (growth function to be precise) at 4 data points. In other words, our procedure takes the hardest slice of VC-dimension and makes an applicable measurement out of it.

The proposed functional complexity measurement is not just implementable *in silico*; it is also experimentally testable. With the high-throughput automated behavior technologies available for larval *Drosophila* [56, 57], one could potentially optogenetically stimulate pairs of input neurons according to patterns illustrated in Figure 1D while recording from the output neurons, such as RGNs and/or descending neurons. It will be exciting to see if the fraction of XORable subnetworks in real larvae match the results obtained via EM-constrained simulations presented here.

An important future direction is to extend this analysis to other EM connectomes [7–11], exploring whether the E-I ratios of other species and in other contexts can also be explained by optimizing for high functional complexity. A comparative study across species would also be of interest. For example, it would be interesting to see if the proposed functional complexity measurement can explain for the expanded inhibitory subnetwork in humans compared to rodents [58]. Meanwhile, a macroscopic gradient of synaptic excitation and inhibition has been observed along the hierarchy of brain regions [59], making it interesting to see if there is also a gradient of functional complexity along the hierarchy of processing.

## 4 Methods

XORable subnetwork Consider a subnetwork defined by two input neurons *I*_1_*, I*_2_ and one output neuron *O*. It includes all possible paths connected between them. We are interested in checking if the binary classification problem XOR, depicted in Figure 1C, is readily solvable by the subnetwork {*I*_1_*, I*_2_*, O*}. To test this, four independent simulations are run (Figure 1D):

1. Simulation 1: only input neuron *I*_2_ is activated at time zero;
2. Simulation 2: both input neurons are activated at time zero;
3. Simulation 3: both input neurons are *not* activated;
4. Simulation 4: only input neuron *I*_1_ is activated at time zero.

The activity of the output neuron *O* is recorded for all time steps. At each times step, we compare the output activity of the four simulations (Figure 1D right panel) and their relative order determines whether the subnetwork is XORable. Specifically, the desired XORable order request simulations 2 & 3 (both input activated or not) to be linearly separable from simulations 1 & 4 (only one of the input activated). Since simulation 3 receives no input activation thus always having zero output, there are only two XORable ordering out of the six possible orderings in total (1 & 4 interchangeable in the illustration). As long as there is at least one time step that have XORable output order then we say the subnetwork is XORable.

Functional Complexity

The functional complexity measurement counts the fraction of input pair (*I*_1_*, I*_2_) that have a robust number (⩾ 20) of XORable subnetworks associated with it, i.e. 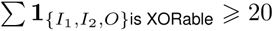.

### Whole-brain larval *Drosophila* EM connectome-constrained RNN

The whole-brain neuronal activity of larval *Drosophila* is simulated with a biologically-constrained recurrent neural network (RNN). Concretely, the neurons’ firing rates **r** at time *t* are given by

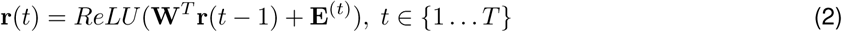

where **W** is the weight matrix describing the connections from pre-synaptic neurons to post-synaptic neurons, with the signs (E-I) sampled and the magnitudes given by the axon-dendrite synaptic count from the EM dataset [3] (all-all synaptic count results are given in supplementaries and are always consistent with the axon-dendrite results). **E**^(^*^t^*^)^ denotes the time-series external input delivered to the input neurons. All simulations were run for

*T* = 20 time steps, a duration long enough for the network dynamics to stabilize when single input neurons were initially activated.

### Uniform sampling on E-I identities

For a target E-I ratio *p*, where *p* ∈ [0, 1], the E-I identity of each neuron is independently sampled from a Bernoulli distribution of probability *p*, which gives the signs of the columns of the weight matrix **W**.

### Connectedness-dependent sampling on E-I identities

For each neuron with a connectedness of *d* (either degree, in degree, or out degree), its E-I identity is sampled from a Bernoulli distribution of probability *p_d_*. The connectedness-dependent probability *p_d_*is set as following: all neurons are grouped into deciles (*q* ∈ [10]) based on their connectedness, and each decile is assigned a probability *p_d_*, forming a probability vector of length 10. In section 2.2, the probability vector is set following exponential function, i.e. *p_d_* ∝ *β^q^*, and the base *β* controls the rate of biasness (Figure 2A). A linear relationship was also explored(Figure A.4), where *p_d_* ∝ *β* × *q* with *β* controlling the rate of biasness. In section 2.3, more connectedness-dependent probability vectors are explored by sampling non-monotone relationships. The most exhaustive way would be search in the [0, 1]^1^0 space for all possible probability vectors, which is computationally prohibitive. Instead, we explored a subset of all length-10 probability vectors featuring various peak positions. Specifically, we generated single-peak probability vectors where the first, fifth and last elements take values from [0, 0.25, 0.5, 0.75, 1], [0, 0.1, 0.2, 0.3, 0.4, 0.5, 0.6, 0.7, 0.8, 0.9, 1], and [0, 0.25, 0.5, 0.75, 1] respectively. The fifth element is constrained to be the extrema, resulting in 184 distinct probability vectors. In addition, three other groups of single-peak vectors were created by shifting the peak two deciles higher, two deciles lower, or five deciles.

### Real brain E-I ratios

The scRNA-seq data were obtained from [13] where 13 individuals were collected for their brains, ventral nerve cord (VNC) or the entire central nervous system (CNS = brain + VNC). Cholinergic cells and Kenyon cells were considered as excitatory. For CNS and VNC samples, only GABAergic cells were considered as inhibitory. For brain samples, glutamatergic cells were additionally considered to be inhibitory.

### E-I prediction low prediction confidence shuffle

In unpublished work, the whole-brain EM connectome neurotransmitter identities are predicted from known subset, leveraging machine learning method [41]. The neurotransmitter prediction of each neuron is associated with a set of confidence scores of the 3 neurotransmitter choices: Acetylcholine, GABAergic, Glutamatergic. Neurons with the maximum confidence score < 0.4 are shuffled for their E-I identities. The shuffling was done 20 times independently.

### Adult *Drosophila* whole-brain EM connectome analysis

All analysis procedures follow larval *Drosophila*, with the following adaptations. The synaptic counts and neurotransmitter predictions are accessed from FlyWire v783 [4], which determines the connection strengths and signs respectively. Only the largest connected component in the constructed whole-brain graph is included in the analysis (132,490 neurons in total with 118,534 neurons with neurotransmitter predictions). All connection strengths are normalized to the percentage of total input onto a given target neuron.

Since the adult *Drosophila* network is two magnitudes larger than the larvae, we had to sub-sample in our functional complexity analysis. Instead of exhaustively iterate through all input combinations as for the larvae, we sub-sampled 10,000 input combinations while still exhaustively iterate through all output neurons. All simulations were run for *T* = 50 time steps, a duration long enough for the network dynamics to stabilize when single input neurons were initially activated.

We also performed a smaller set of connectedness-dependent E-I sampling but with the same coverage on E-I ratio and differential connectedness. Specifically, we sampled from 235 different connectedness-dependent probability vectors, covering uniform (0.1, 0.2, 0.3, 0.5, 0.7, 0.8, 0.9), exponential (28 probability vectors), and non-monotonic relationships (200 probability vectors).

### Sparse network training

All networks are trained with stochastic gradient descent on Fashion-MNIST with 7000 training examples and 10,000 validation examples. The learning rate and batch size are selected based on grid search and chosen based on the best average across all scenarios. The sparsification and signs are initialized based on settings specified in Table 2. During training, the wiring and E-I identity of each connection is allowed to be updated in classical pruning only, for all other 3 scenarios following Dale’s rule, the zero mask and signs are fixed throughout training.

**Table 1:** All possible orders of output responses of the four simulations at each time step. (related to Figure 1)

| Relative order of the four simulations' output neuron response | XORable? |
| --- | --- |
| $3 < 1 < 2 < 4$ | n |
| $3 < 1 < 4 < 2$ | n |
| $3 < 2 < 1 < 4$ | y |
| $3 < 2 < 4 < 1$ | y |
| $3 < 4 < 1 < 2$ | n |
| $3 < 4 < 2 < 1$ | n |

**Table 2:** Sparse network E-I configuration specifications across scenarios. (related to Figure 5)

| E-I scenario as in Figure 5 | E(E+I) | Exc sparsity | Inhib sparsity | Overall sparsity | $(w_{>0})/(w_{>0} + w_{<0})$ |
| --- | --- | --- | --- | --- | --- |
| 80% Exc (Dale's) | 0.8 | 0.99375 | 0.975 | 0.99 | 0.5 |
| 50% Exc (Dale's) | 0.5 | 0.99 | 0.99 | 0.99 | 0.5 |
| 20% Exc (Dale's) | 0.2 | 0.975 | 0.99375 | 0.99 | 0.5 |
| classical pruning | N/A | 0.99 | 0.99 | 0.99 | 0.5 |

## Acknowledgements

This work was supported by NSF NeuroNex Award (Grant no. 2014862) (to J.V.), NSF CAREER Award (Grant no. 1942963) (to J.V.), the JHU MINDS Fellowship (to Q.W.), and the Francis Crick Institute core funding (to M.W.), including Cancer Research UK (CC2252), the UK Medical Research Council (CC2252), and the Wellcome Trust (CC2252).

## Author contributions

Conceptualization: Q.W.,C.E.P.,and J.T.V. Methodology: Q.W. and C.E.P. Analysis: Q.W. DNN training: Q.W. Larva *Drosophila* neurotransmitter prediction: N.R., Y.Y., C.S., A.S., M.W., A.C., and M.Z. Supervision: C.E.P. and J.T.V. Writing - original draft: Q.W. Writing - Review and editing: Q.W., J.T.V., C.E.P., M.W., M.Z., and A.C.

## A Supplementary Figures

**Figure A.1:**
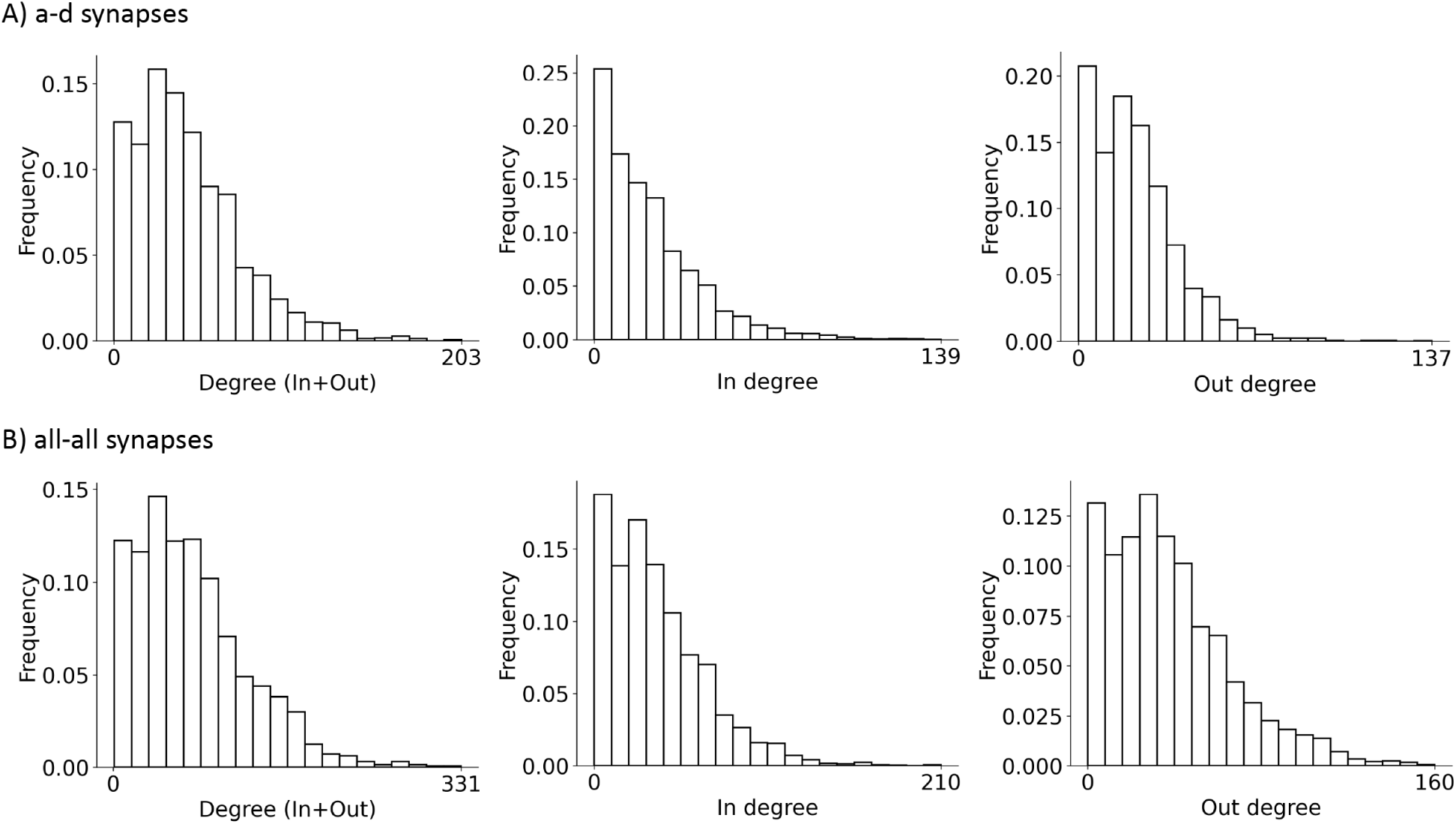
The degree distribution of larval *Drosophila* EM connectome. A) Connections given by axon-dendritic synapses; B) Connections given by all four types of synapses.

**Figure A.2:**
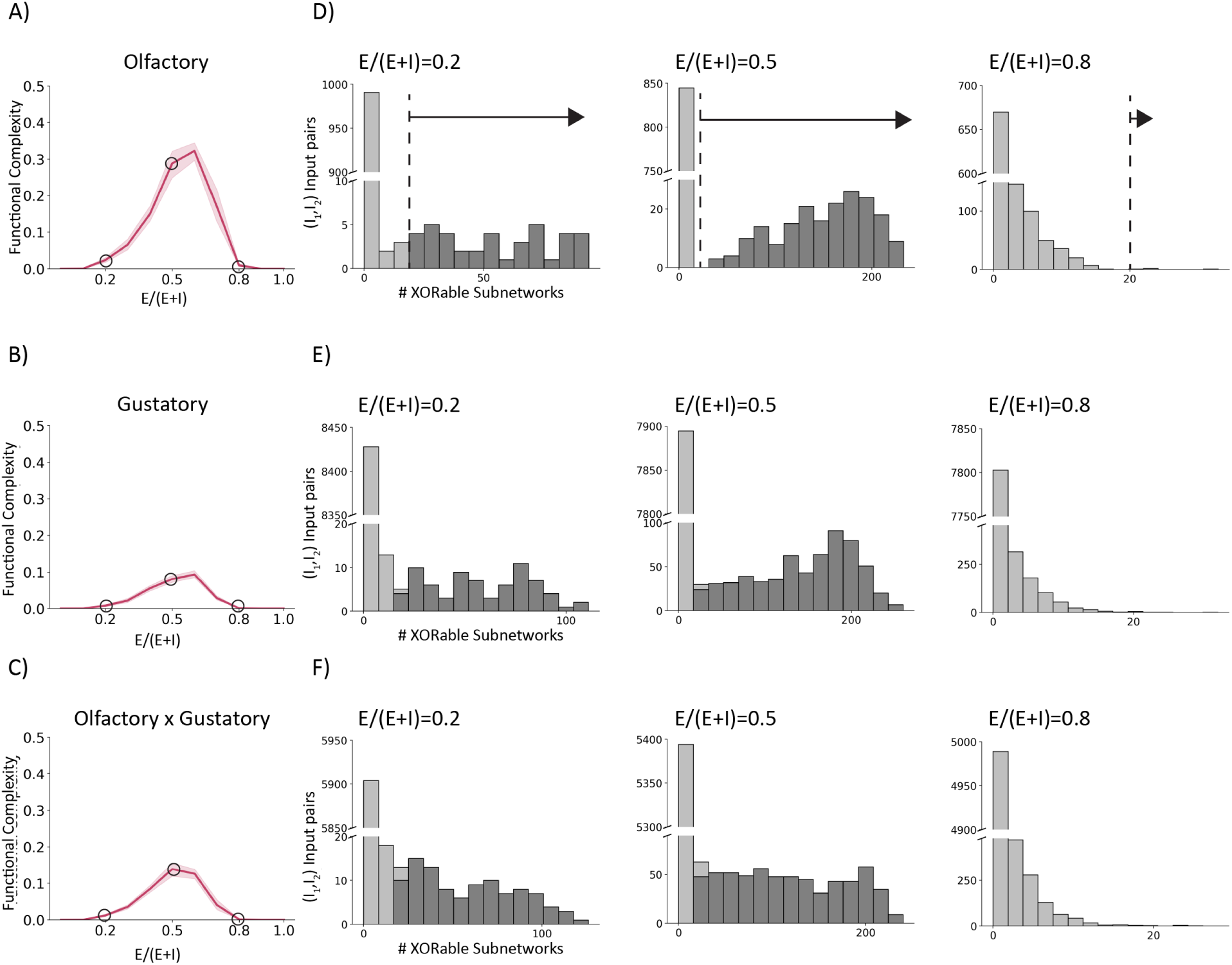
Distribution of XORable subnetworks when E-I identities are uniformly sampled across the entire network. (related to Figure 2) A-C) Functional complexity of networks with different E-I ratios (E-I identities sampled uniformly for all neurons in the EM-constrained RNN). The functional complexities are measured with subnetworks receiving inputs from different sensory modalities, {*I*1*, I*2} =: A) {Olfactory_1_, Olfactory_2_}, B) {Gustatory_1_, Gustatory_2_}, C) {Olfactory, Gustatory}. Shaded area mark the 95% confidence interval (n=10). Circles mark the corresponding points shown in D-F on the right. D-F) Distributions of XORable subnetworks grouped by the {*I*1*, I*2} input pairs. Each histogram is of a particular sensory modality (given on the left), and of a single RNN with E-I identities sampled at the ratio indicated on top. The functional complexity is then calculated by the fraction of XORable subnetworks, excluding the ones where a unique {*I*1*, I*2} input pair have less than 20 output neurons exhibitng XORable pattern (out of 400 in total) (details in Sec 4). Subnetworks counted towards functional complexity are marked by dark gray.

**Figure A.3:**
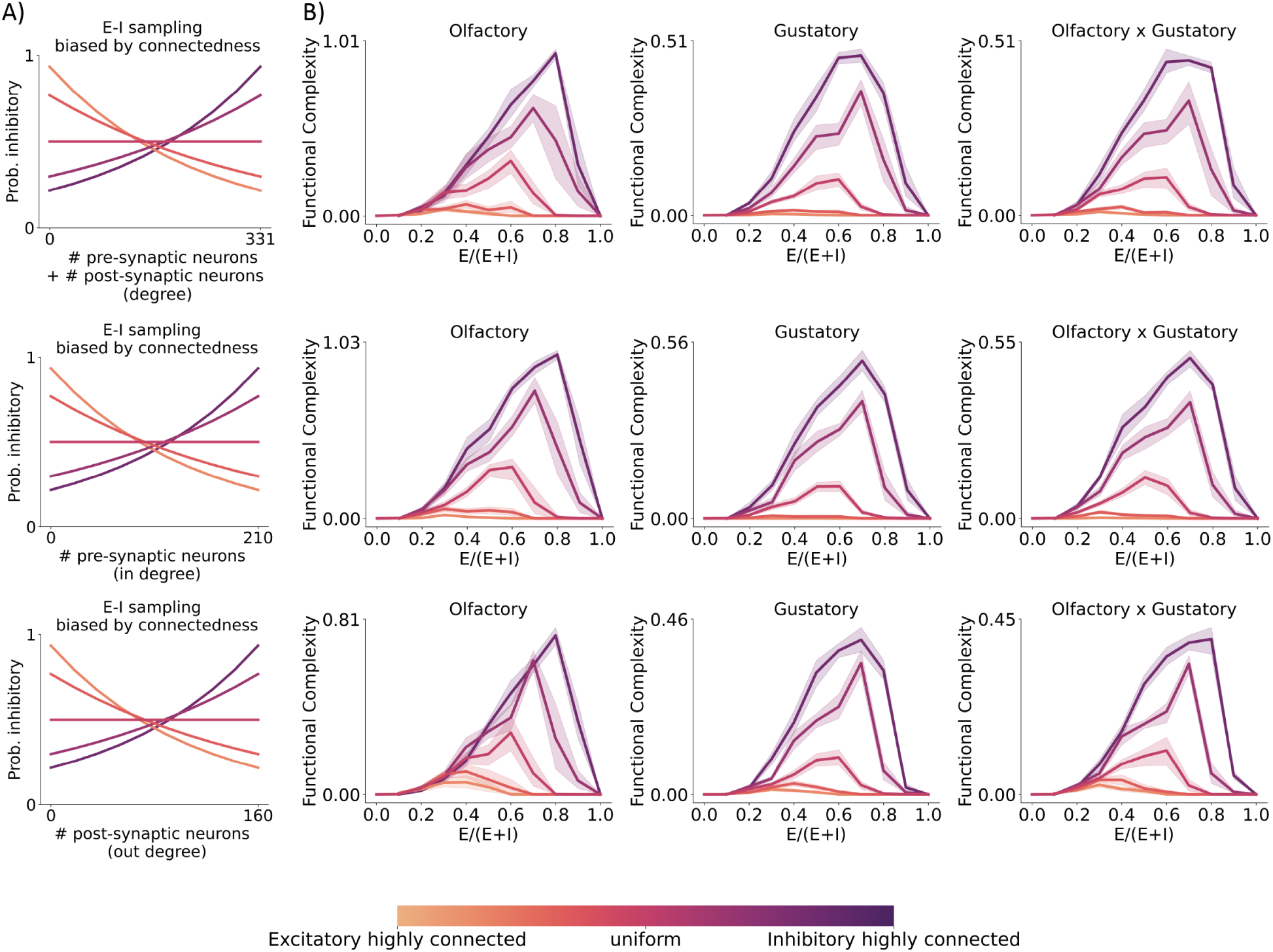
Same as Figure 2 except including all-to-all synaptic counts as the EM constraint.

**Figure A.4:**
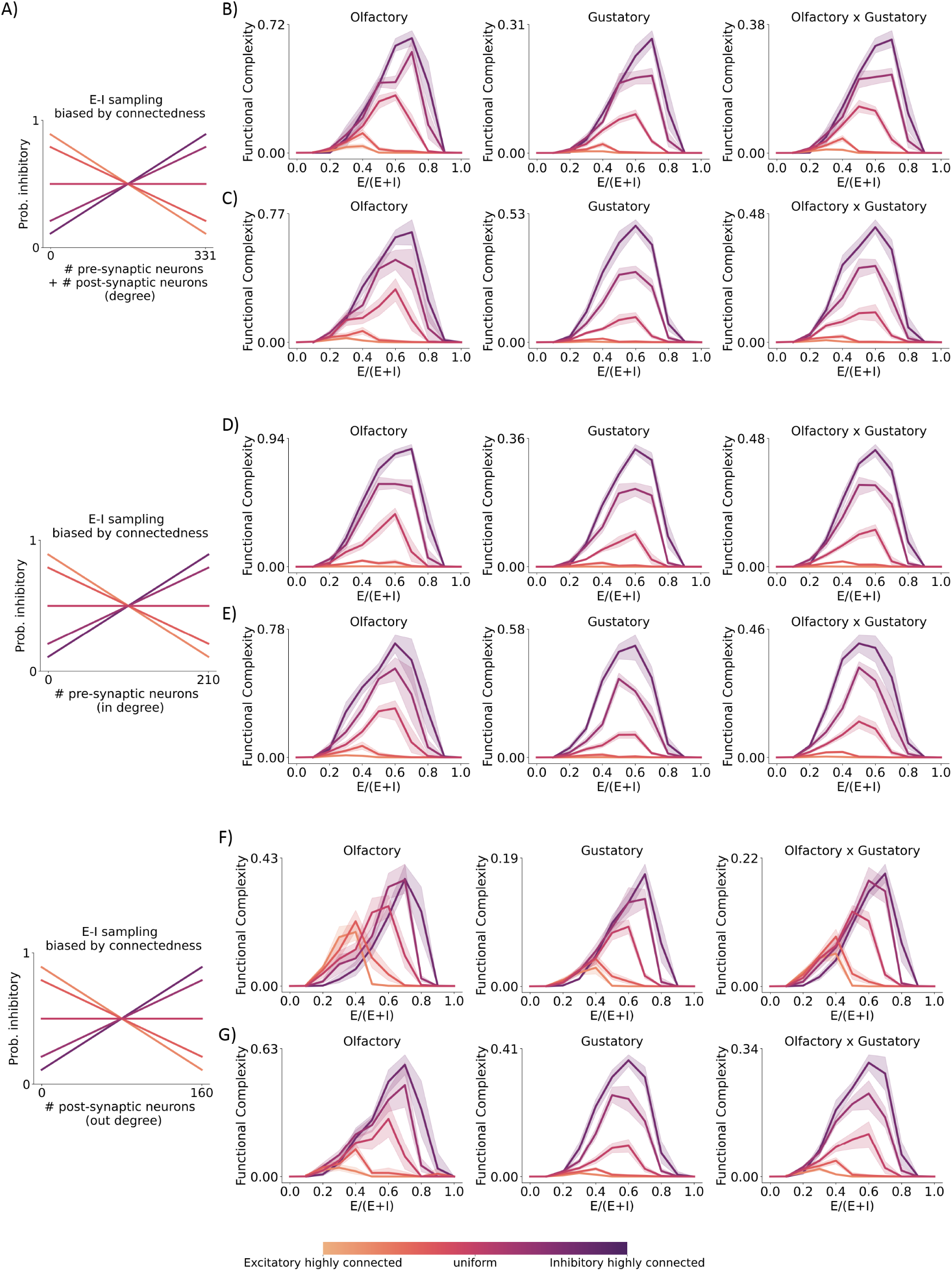
Same as Figure 2 except for linear degree-dependent sampling procedures. A) Linear degree-dependent E-I probability sampling curves. B,D,F) Results for axon-dendrite synapses as the EM constraint. C,E,G) Results for all-to-all synapses as the EM constraint.

**Figure A.5:**
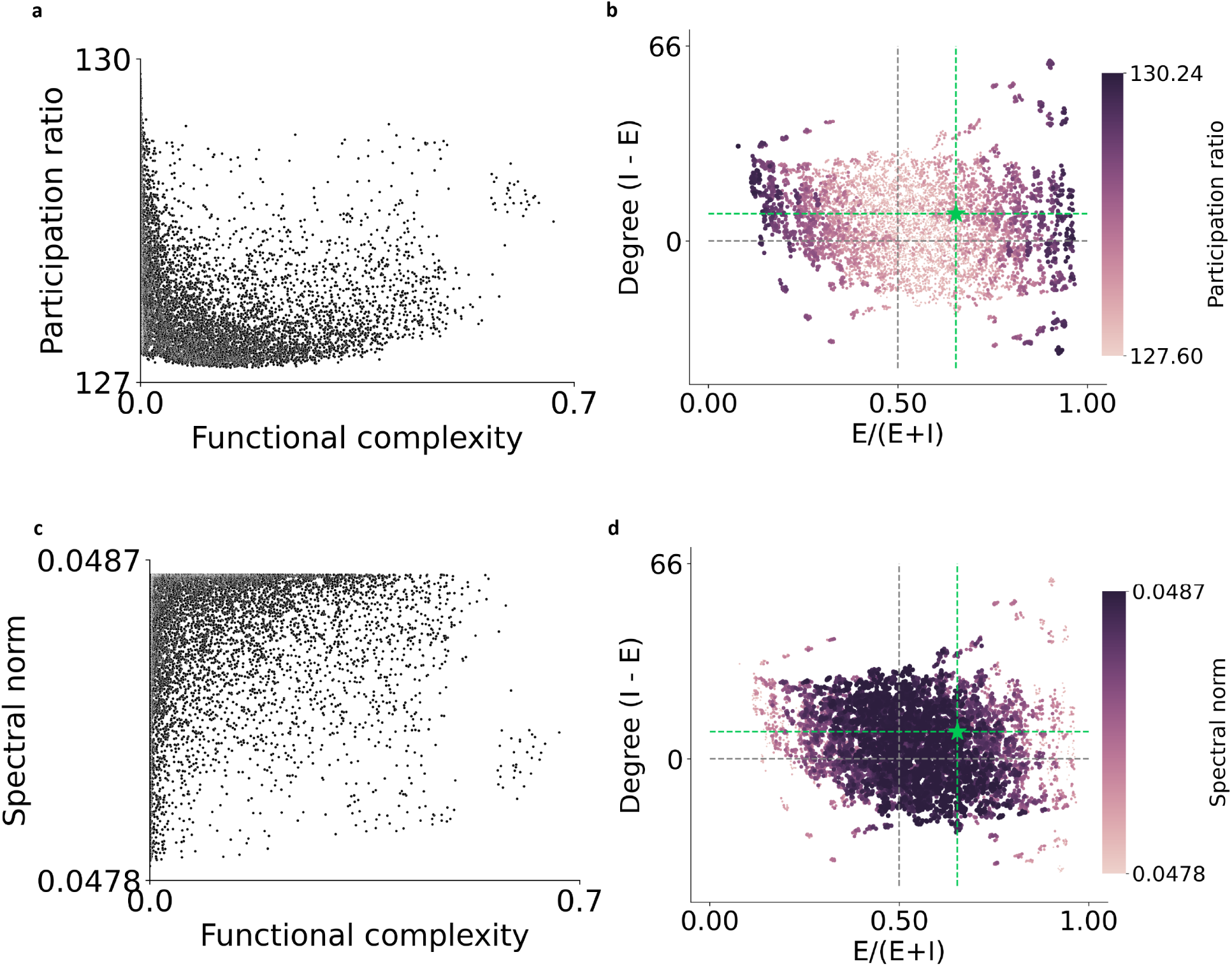
Other methods of complexity measurements cannot explain for the network statistics observed in the brain. **a**, For all sampled networks based on the larval *Drosophila* EM connectome, we compared its participation ratio, which quantifies the effective dimension of the matrix, to our proposed functional complexity measurement. Evidently, many networks with low functional complexity have a high companion participation ratio. **b**, the high participation ratio networks prefer all excitatory (or inhibitory) compositions and do not show a difference across different degree distributions (y-axis). The high participation ratio networks do not align with the real brain observation (green star). **c**, comparison of our functional complexity measurement to spectral norm which shows little correlation. **d**, the spectral norm of the sampled networks show little variation (notice the small range of the color bar).

**Figure A.6:**
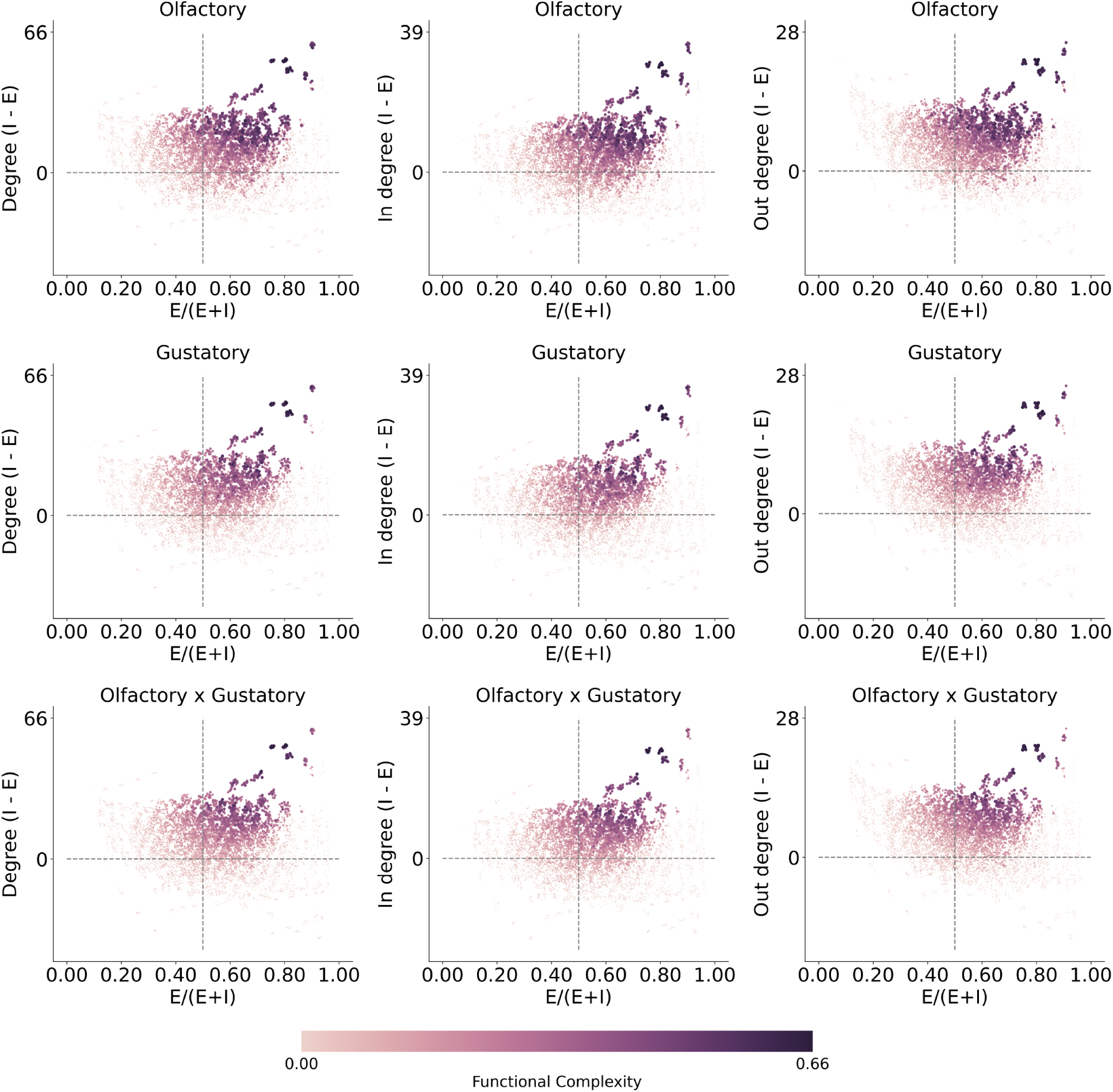
Related to Figure 3, functional complexity plotted separately across sensory modalities. Degree-dependent sampling is based on degree (In+Out). Across sensory modalities (top to bottom) and degree quantification methods (left to right), networks of higher functional complexity (lighter color) have inhibitory neurons more highly connected than excitatory neurons (above horizontal gray line) and there is an over-abundance of excitatory neurons (right to vertical gray line).

**Figure A.7:**
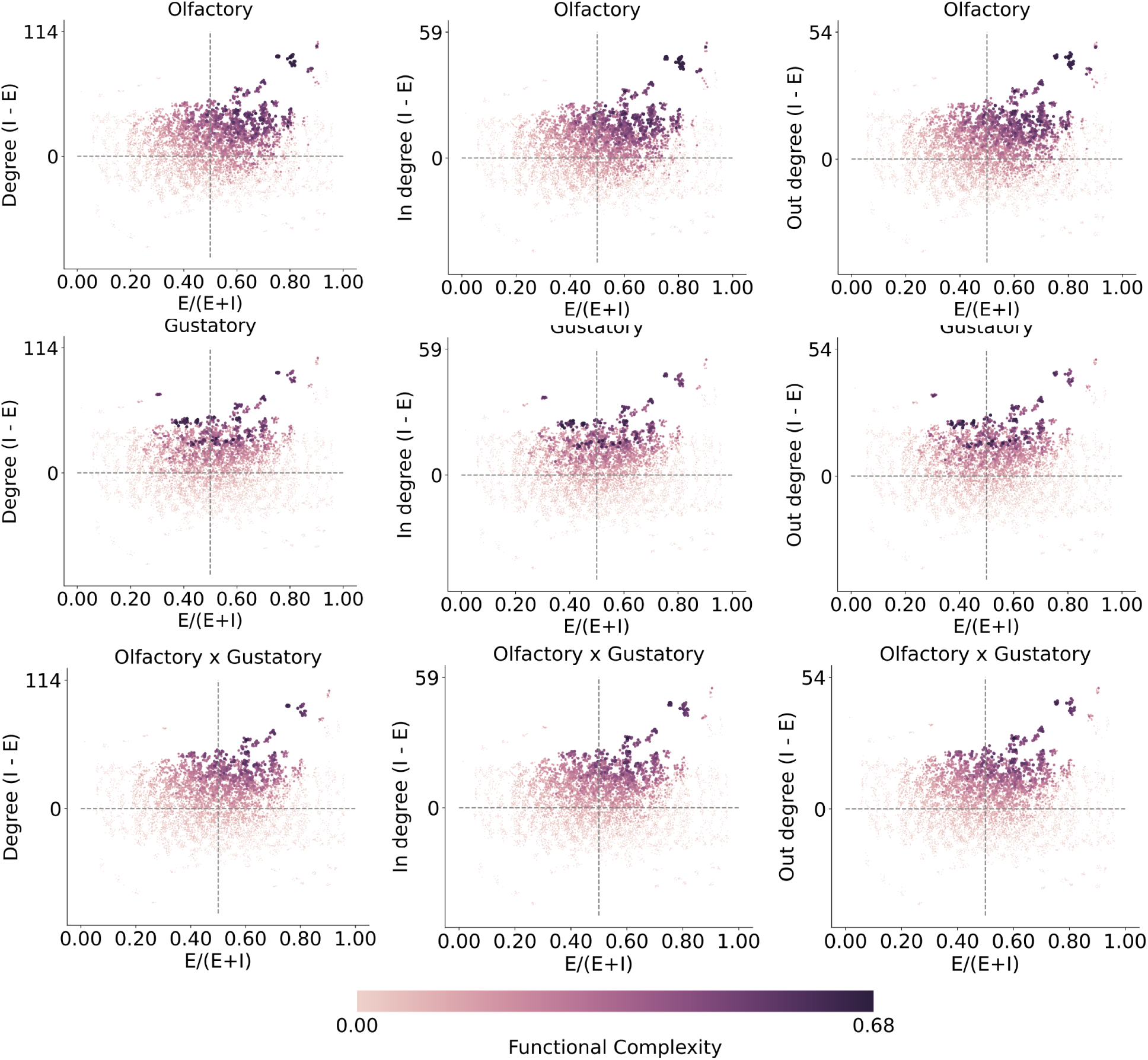
Related to Figure 3, functional complexity plotted separately across sensory modalities. Degree-dependent sampling is based on degree (In+Out) and the connection weight is given by all-all synaptic counts. Across sensory modalities (top to bottom) and degree quantification methods (left to right), networks of higher functional complexity (lighter color) have inhibitory neurons more highly connected than excitatory neurons (above horizontal gray line) and there is an over-abundance of excitatory neurons (right to vertical gray line).

**Figure A.8:**
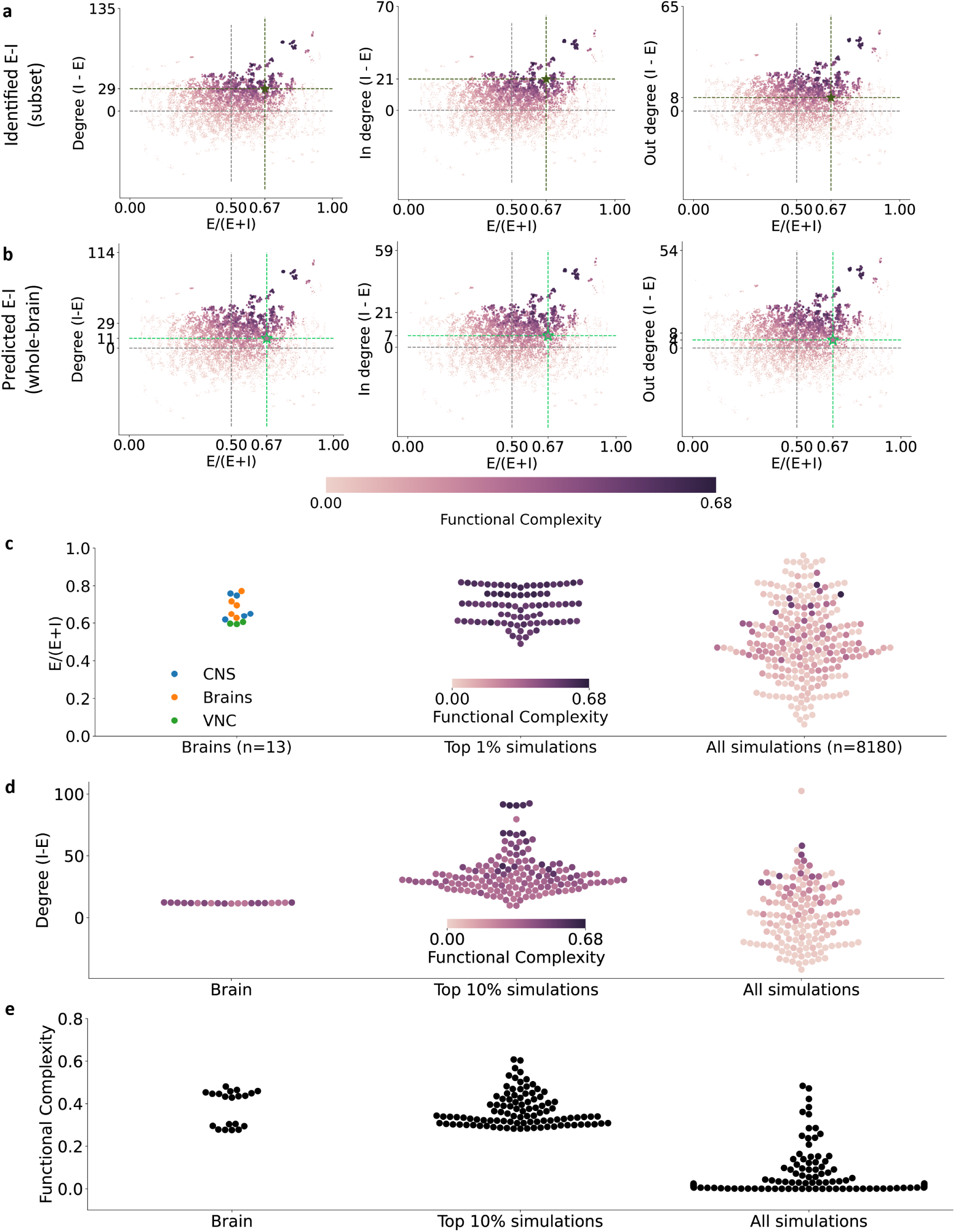
Same as Figure 3 except including all-to-all synaptic counts as the EM constraint. The mean E-I ratio of the top 13 group-averages and top 1% simulations are **not** statistical significantly different from the 13 brain E-I ratios (two-sided two-sample t-test, p-val: 0.99, 0.94 respectively); while the mean of all simulations are statistical significantly different from the brain observations (p-val: 0.01). Multiple comparisons corrected by Dunnett’s.

**Figure A.9:**
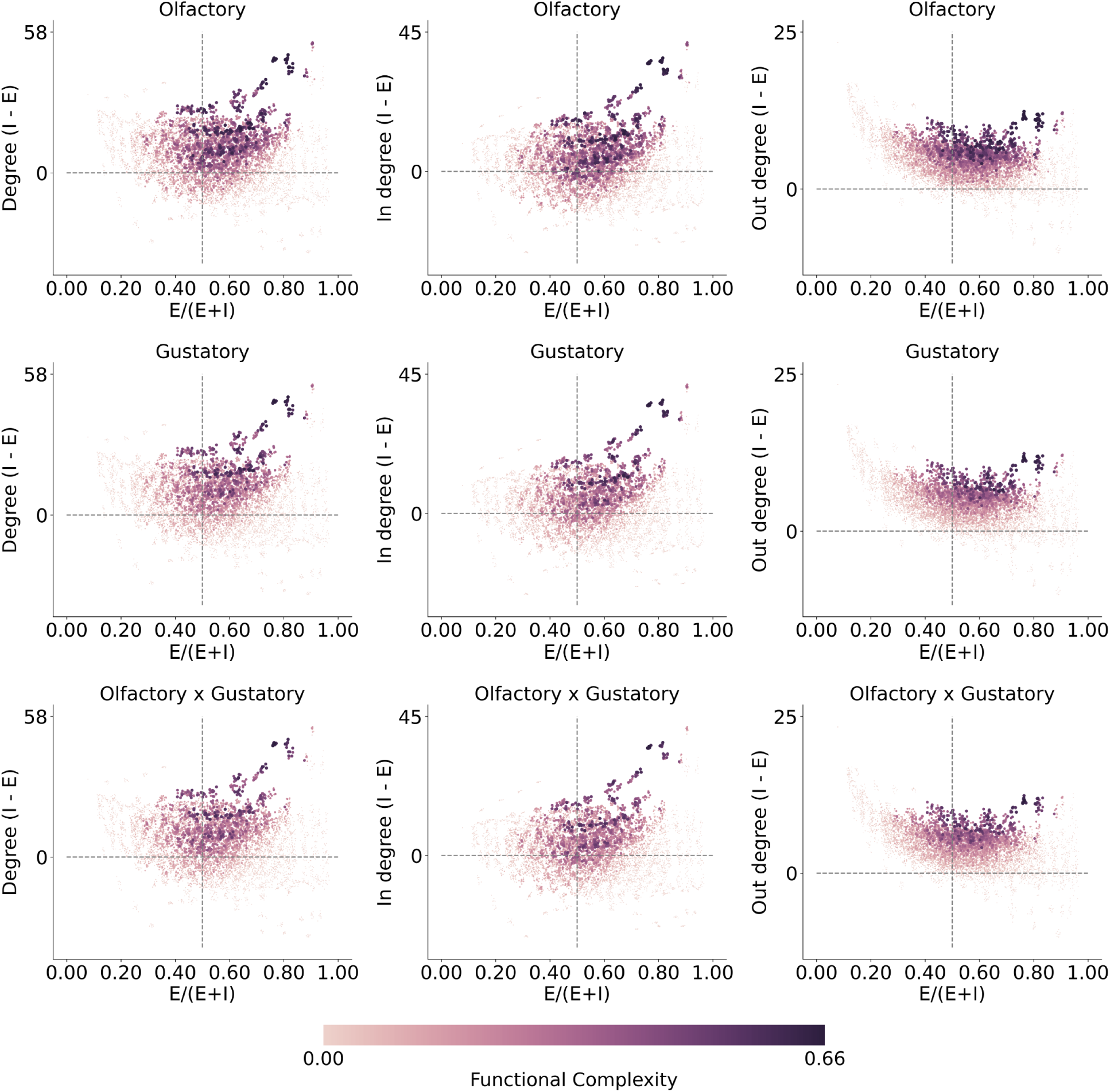
Related to Figure 3, functional complexity plotted separately across sensory modalities. Degree-dependent sampling is based on in degree . Across sensory modalities (top to bottom) and degree quantification methods (left to right), networks of higher functional complexity (lighter color) have inhibitory neurons more highly connected than excitatory neurons (above horizontal gray line) and there is an over-abundance of excitatory neurons (right to vertical gray line).

**Figure A.10:**
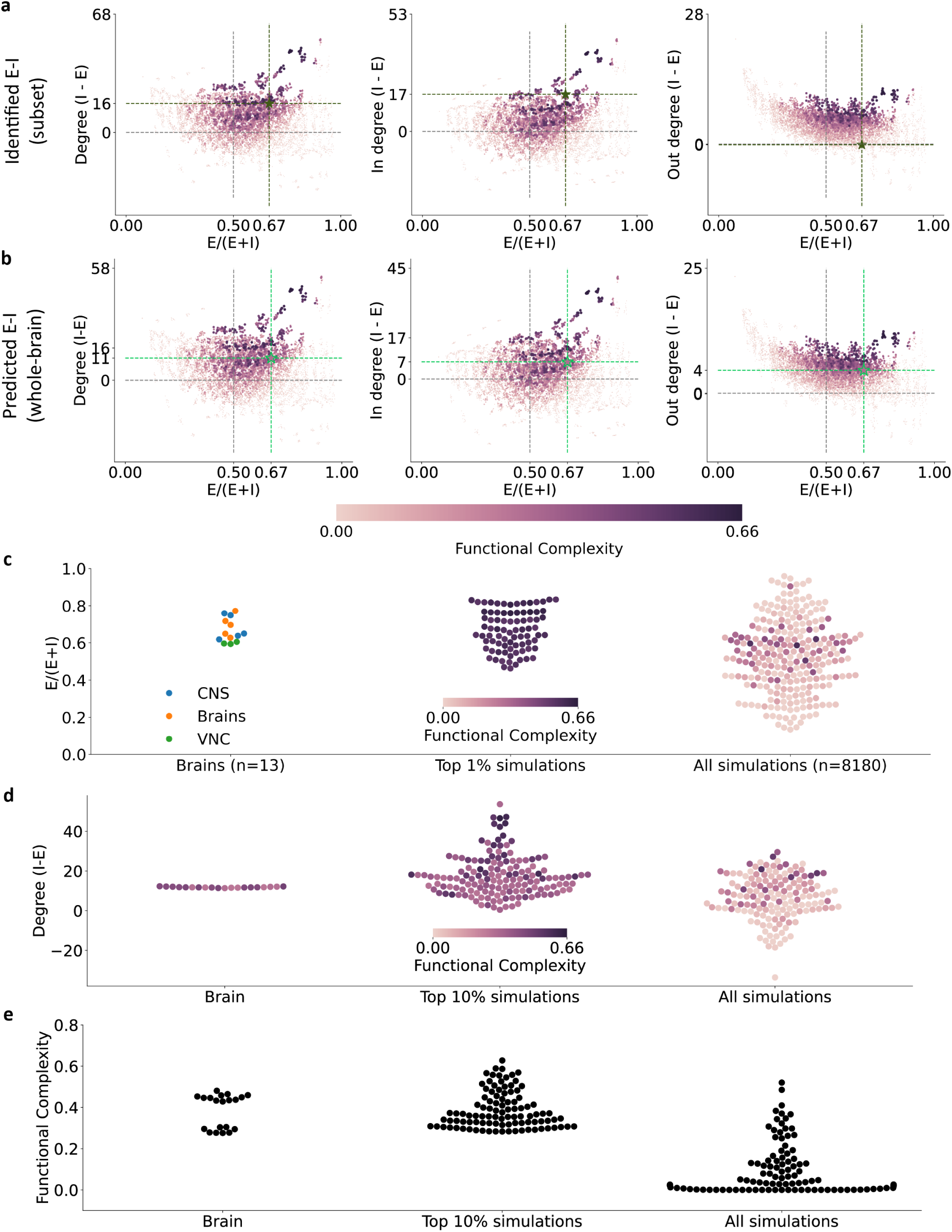
Same as Figure 3 except the degree-dependent sampling is based on in degree. The mean E-I ratio of the top 13 group-averages and top 1% simulations are **not** statistical significantly different from the 13 brain E-I ratios (two-sided two-sample t-test, p-val: 0.97, 0.99 respectively); while the mean of all simulations are statistical significantly different from the brain observations (p-val: 0.03). Multiple comparisons corrected by Dunnett’s.

**Figure A.11:**
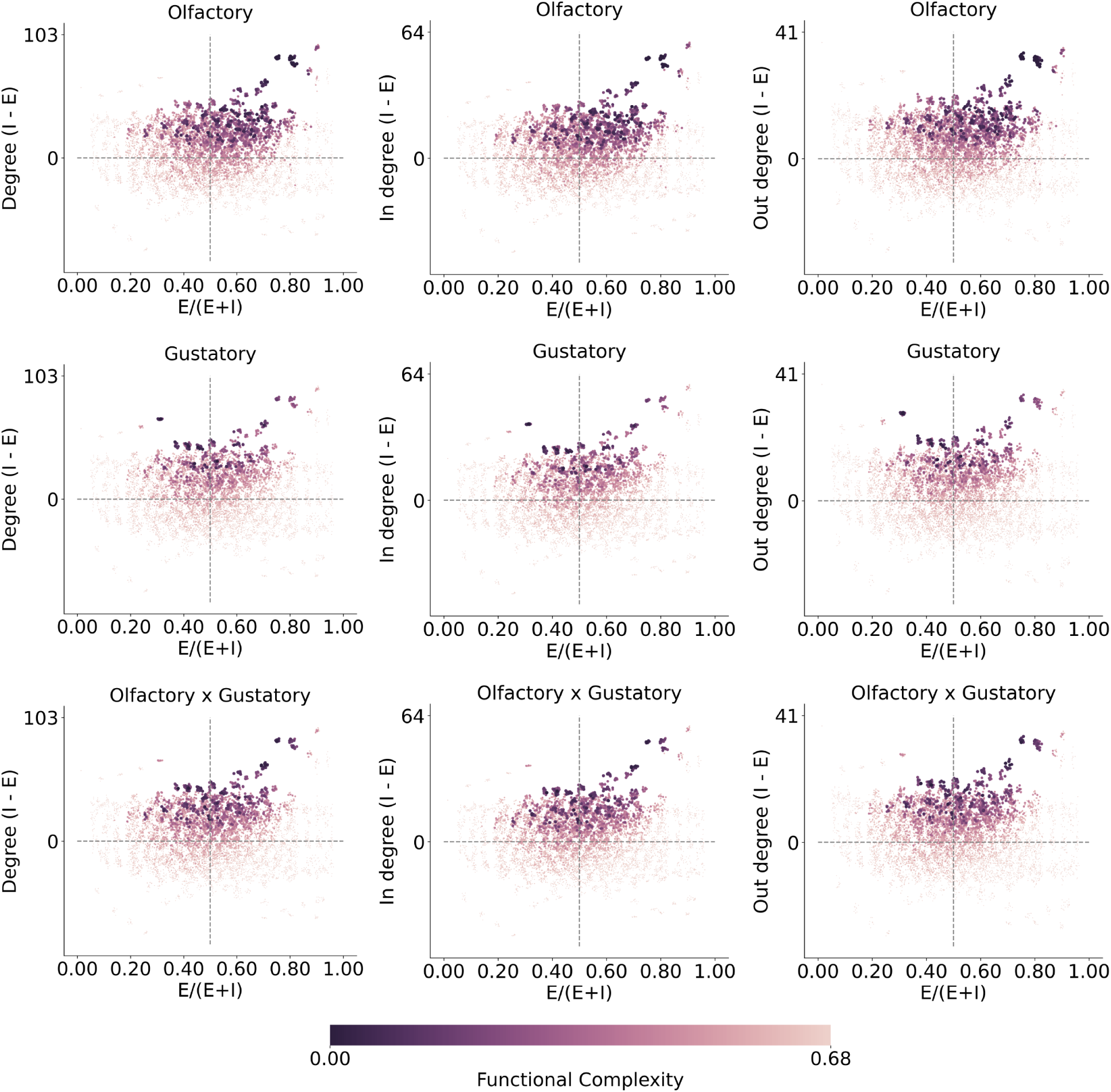
Related to Figure 3, functional complexity plotted separately across sensory modalities. Degree-dependent sampling is based on in degree and the connection weight is given by all-all synaptic counts. Across sensory modalities (top to bottom) and degree quantification methods (left to right), networks of higher functional complexity (lighter color) have inhibitory neurons more highly connected than excitatory neurons (above horizontal gray line) and there is an over-abundance of excitatory neurons (right to vertical gray line).

**Figure A.12:**
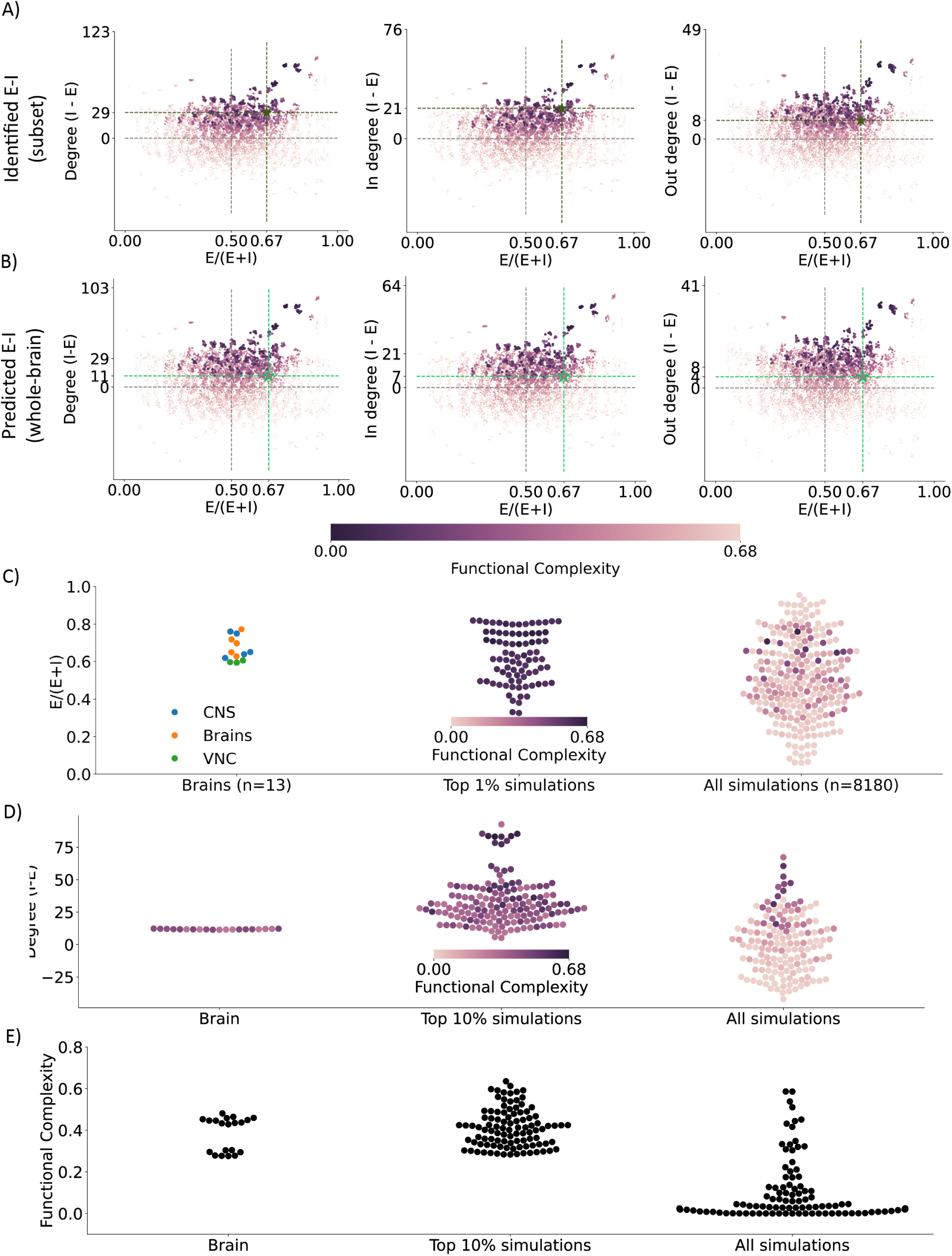
Same as Figure 3 except the degree-dependent sampling is based on in degree and including all-to-all synaptic counts as the EM constraint. The mean E-I ratio of the top 13 group-averages and top 1% simulations are **not** statistical significantly different from the 13 brain E-I ratios (two-sided two-sample t-test, p-val: 0.65, 0.70 respectively); while the mean of all simulations are statistical significantly different_3_f_2_rom the brain observations (p-val: 0.01). Multiple comparisons corrected by Dunnett’s.

**Figure A.13:**
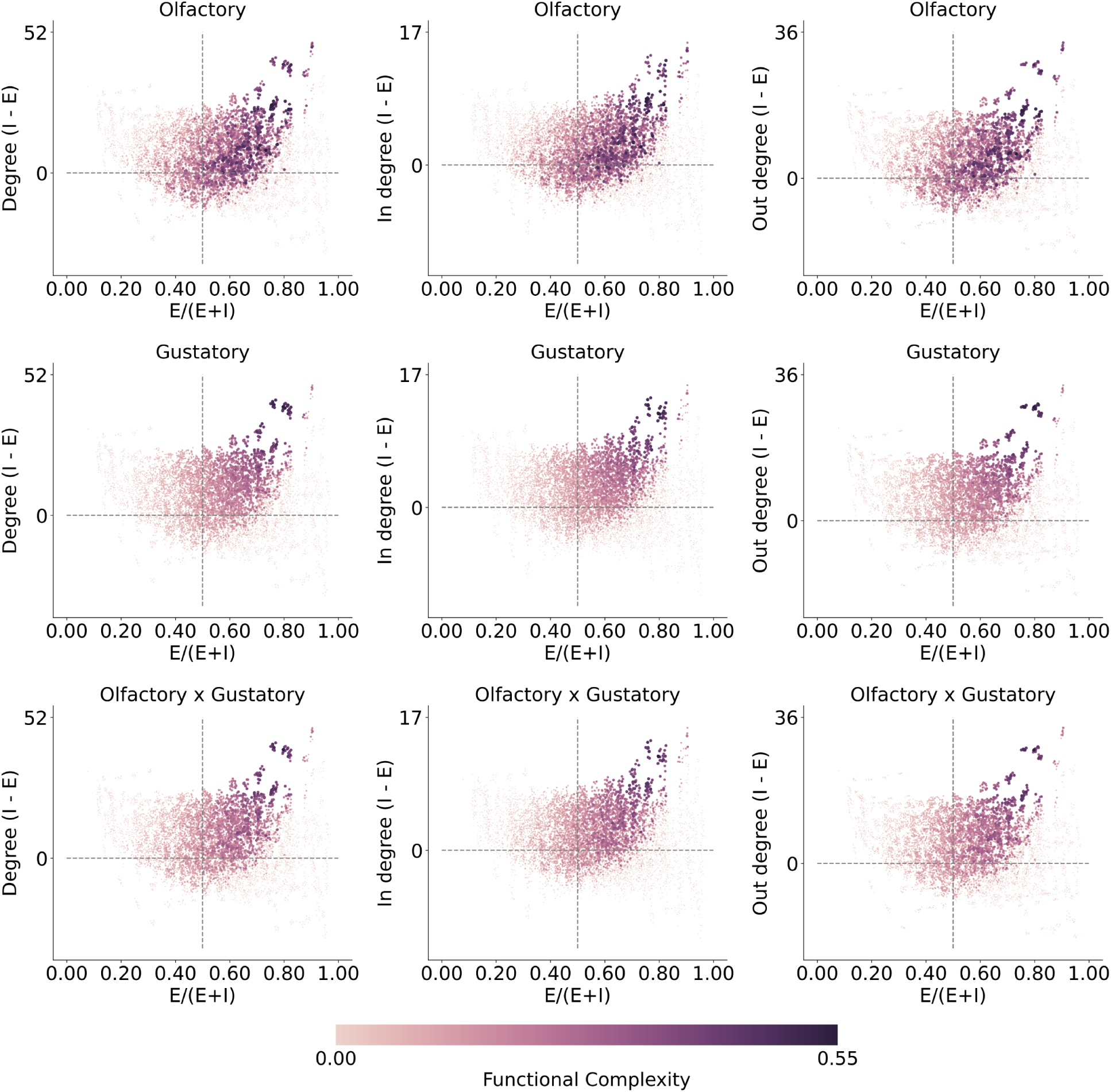
Related to Figure 3, functional complexity plotted separately across sensory modalities. Degree-dependent sampling is based on out degree. Across sensory modalities (top to bottom) and degree quantification methods (left to right), networks of higher functional complexity (lighter color) have inhibitory neurons more highly connected than excitatory neurons (above horizontal gray line) and there is an over-abundance of excitatory neurons (right to vertical gray line).

**Figure A.14:**
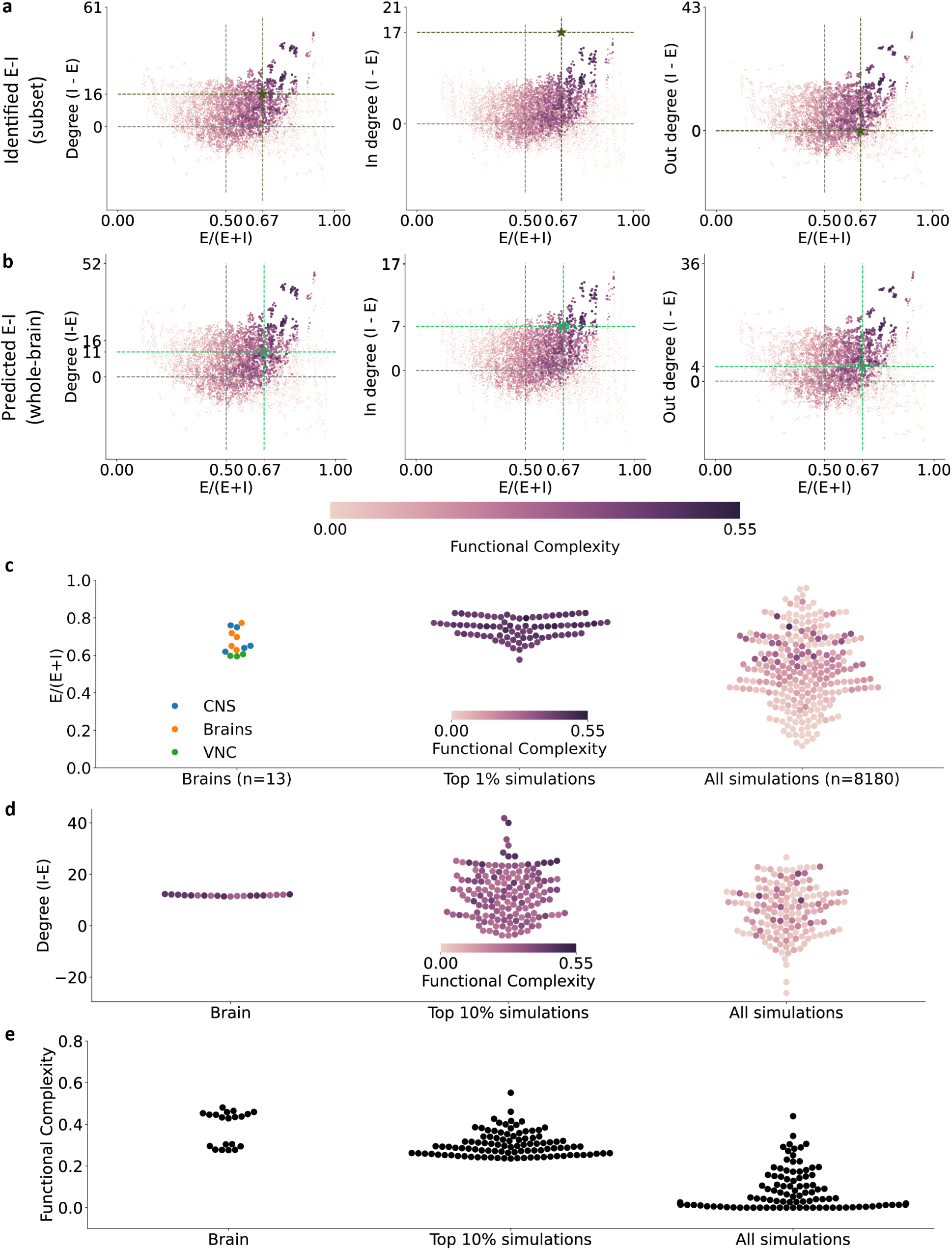
Same as Figure 3 except the degree-dependent sampling is based on out degree and including all-to-all synaptic counts as the EM constraint. The mean E-I ratio of the top 13 group-averages and top 1% simulations are **not** statistical significantly different from the 13 brain E-I ratios (two-sided two-sample t-test, p-val: 0.51, 0.30 respectively); while the mean of all simulations are statistical significantly different_3_f_4_rom the brain observations (p-val: 0.03). Multiple comparisons corrected by Dunnett’s.

**Figure A.15:**
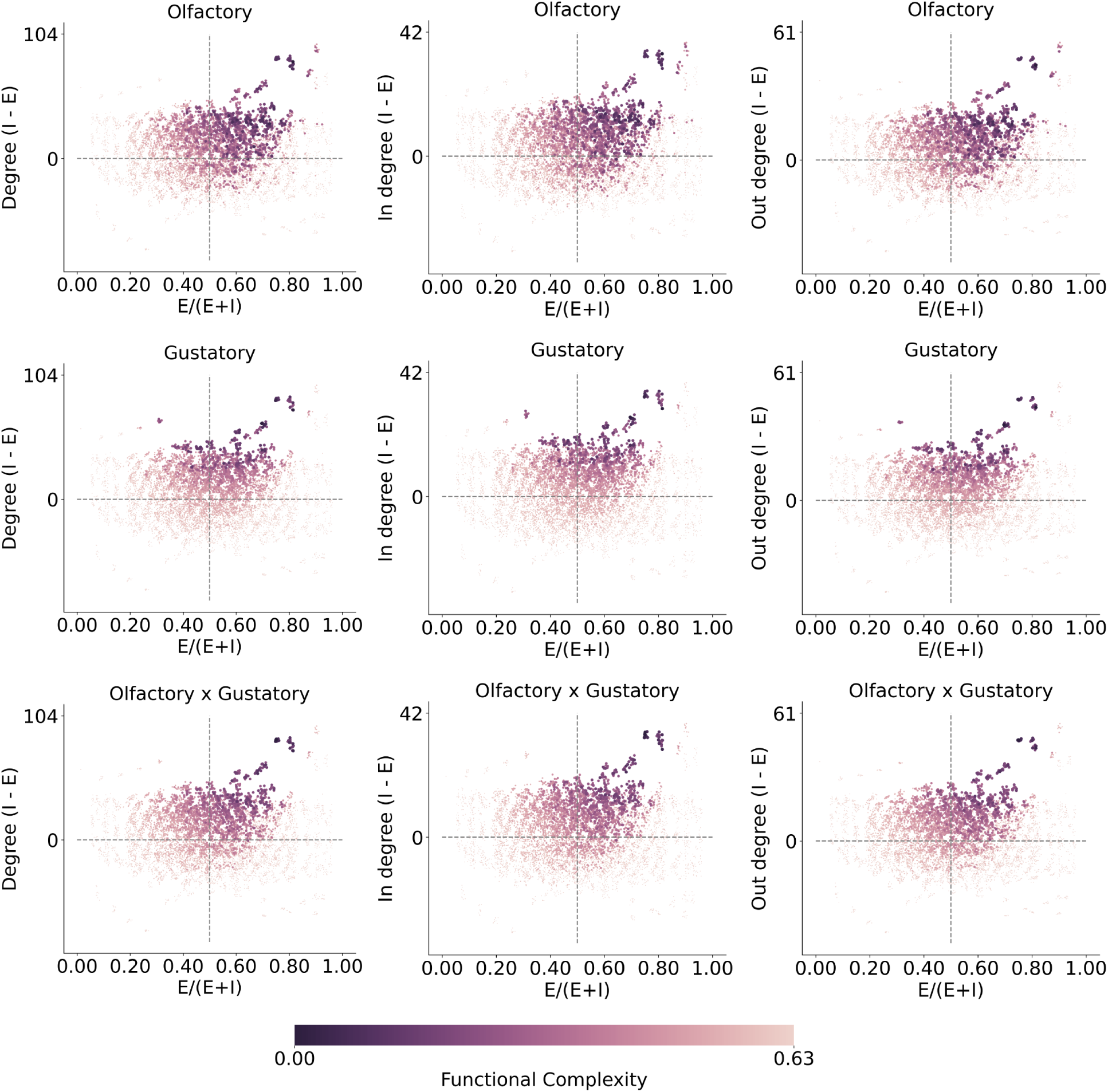
Related to Figure 3, functional complexity plotted separately across sensory modalities. Degree-dependent sampling is based on out degree and the connection weight is given by all-all synaptic counts. Across sensory modalities (top to bottom) and degree quantification methods (left to right), networks of higher functional complexity (lighter color) have inhibitory neurons more highly connected than excitatory neurons (above horizontal gray line) and there is an over-abundance of excitatory neurons (right to vertical gray line).

**Figure A.16:**
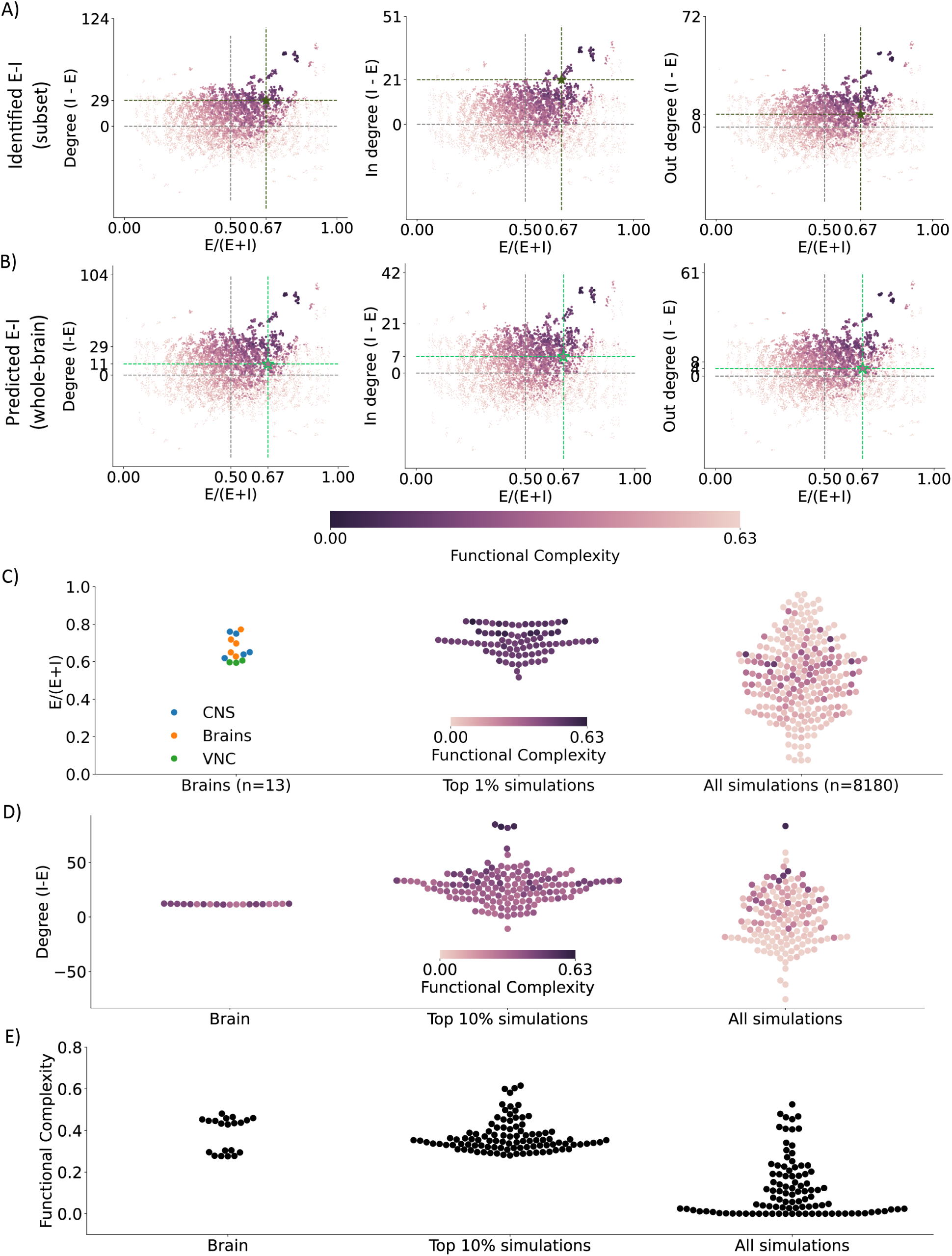
Same as Figure 3 except the degree-dependent sampling is based on out degree and including all-to-all synaptic counts as the EM constraint. The mean E-I ratio of the top 13 group-averages and top 1% simulations are **not** statistical significantly different from the 13 brain E-I ratios (two-sided two-sample t-test, p-val: 0.92, 0.76 respectively); while the mean of all simulations are statistical significantly different3f6rom the brain observations (p-val: 0.01). Multiple comparisons corrected by Dunnett’s.

**Figure A.17:**
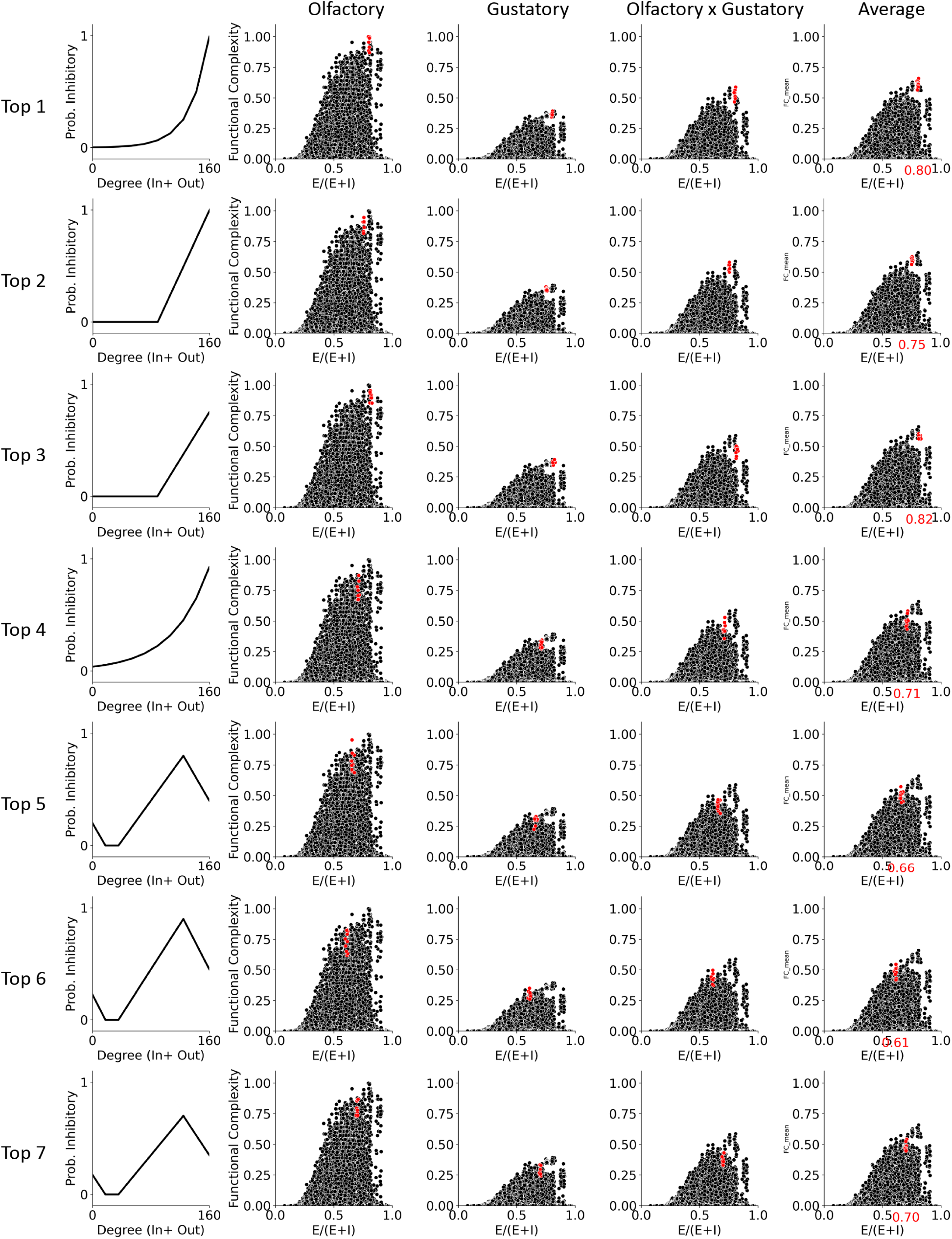
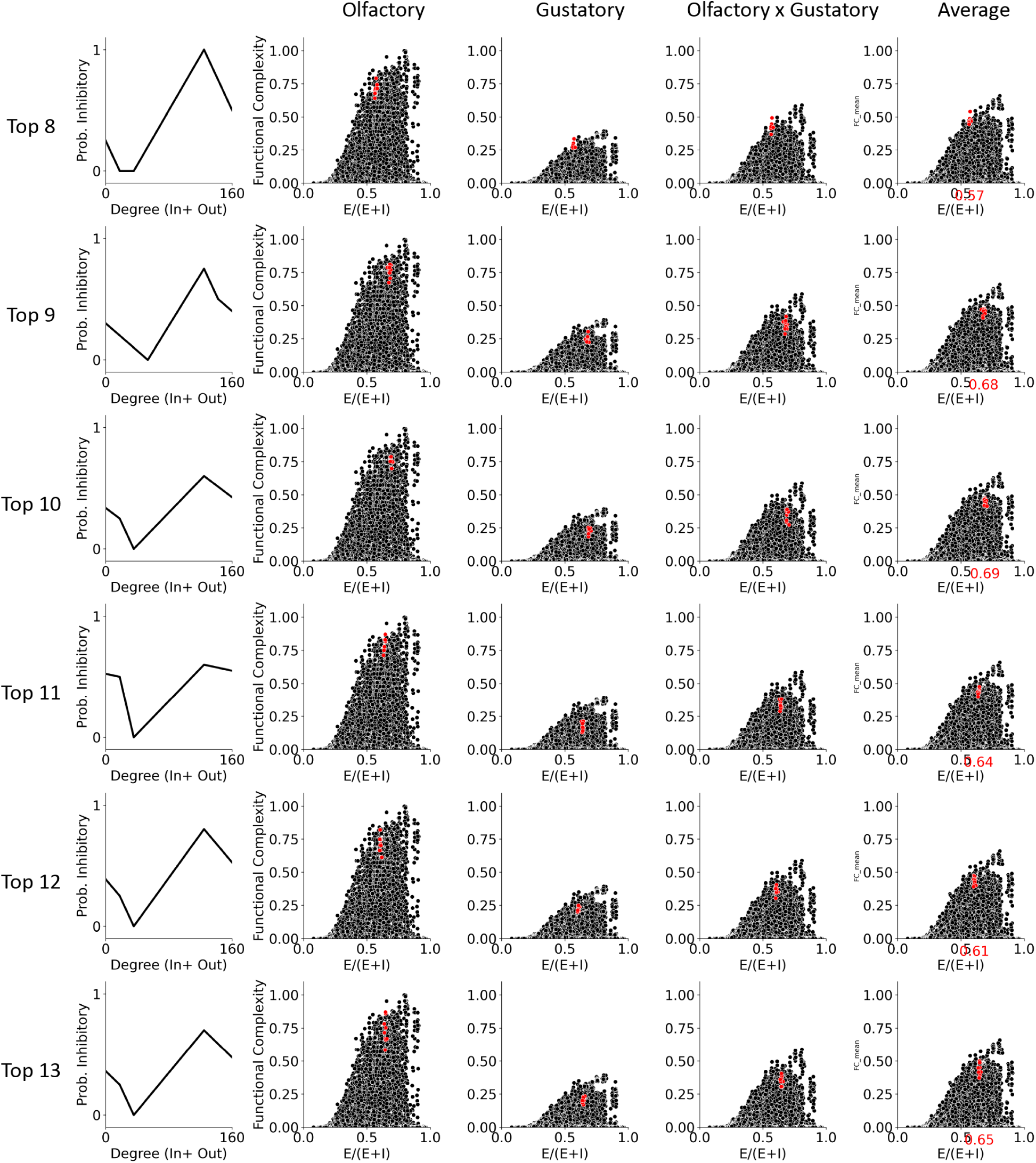
Top 13 degree-dependent E-I probability functions. (Related to Figure 3) The top 13 E-I ratios plotted in Figure 3 middle left are delineated here. Top to bottom: the E-I probability function group that resulted in the highest functional complexity to lower.

**Figure A.18:**
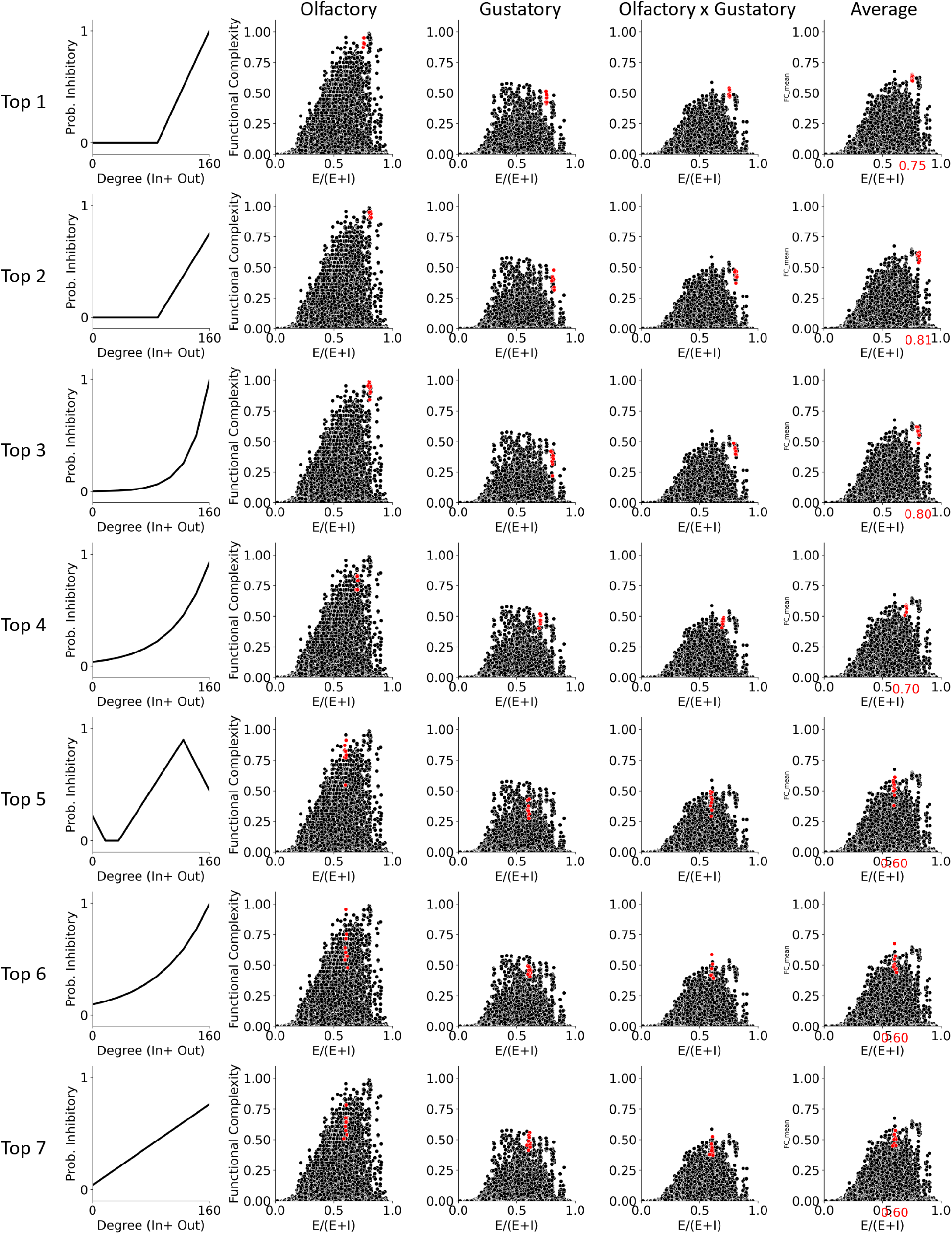
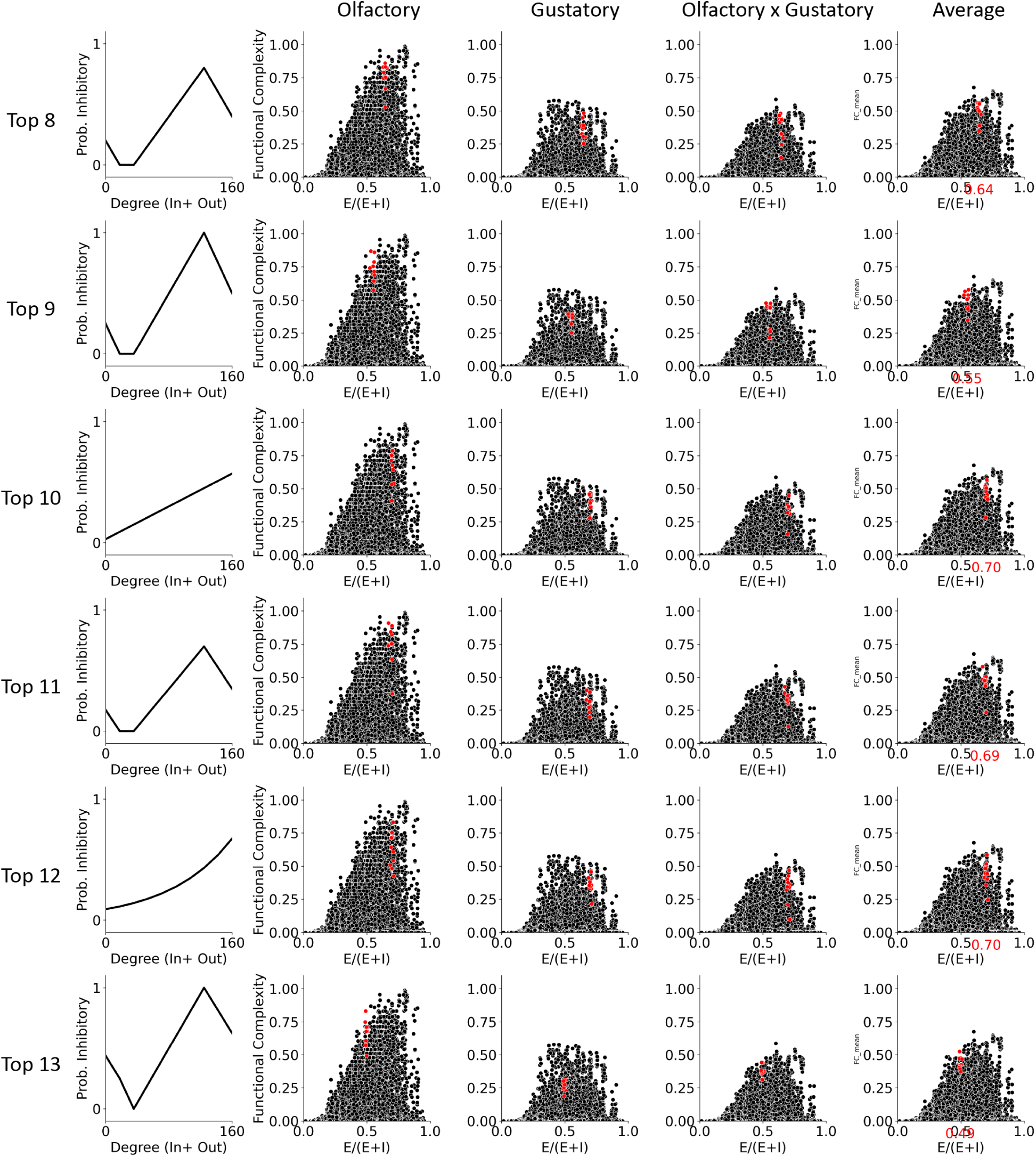
Same as Supplementary Figure A.17 except including all-to-all synaptic counts as the EM constraint.

**Figure A.19:**
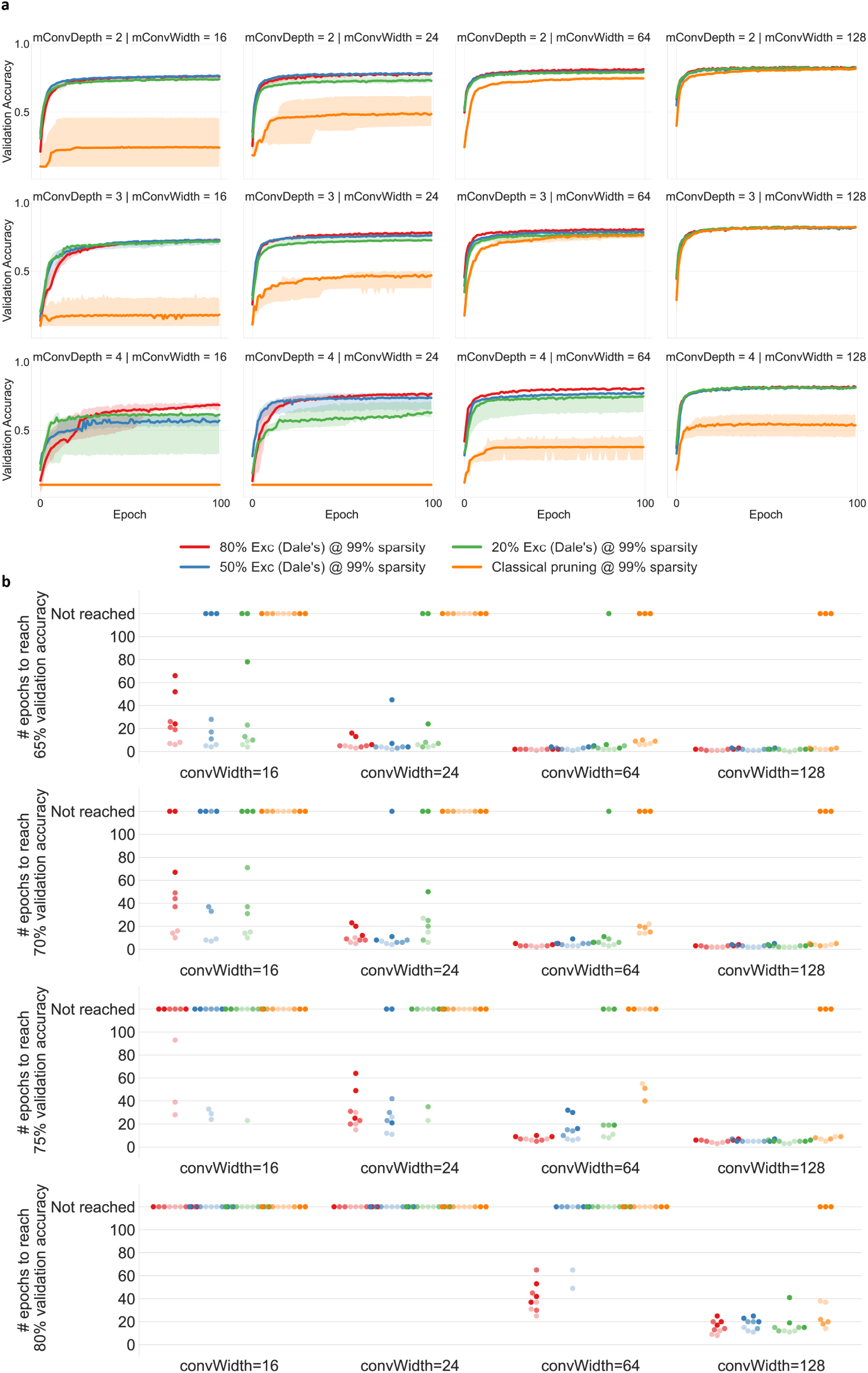
Related to Figure 5. A) Convolutional neural networks (CNNs) with 99% sparsity are trained with unconstained E-I configurations (orange, classical pruning setting), biologically constrained E-I configurations that follow the Dale’s rule with 80% of excitatory neurons (red), 50% excitatory neurons (blue), and 20% excitatory neurons. Under classical pruning procedures (orange), CNNs of various depth (across rows) and width (across columns) can face difficulty in training at 99% sparsity, especially when the networks have narrow widths. These difficulties can be remedied by biologically constrained E-I configurations (blue, red, green). All plots are median with shaded area showing min and max (n=3). B) Computation efficiency

## Footnotes

1 Universal approximation theory is the foundational theoretical guarantee that helped to catapult deep learning out of the first AI winter [31–33].

